# Protein language model powers accurate and fast sequence search for remote homology

**DOI:** 10.1101/2023.04.03.535375

**Authors:** Wei Liu, Ziye Wang, Ronghui You, Chenghan Xie, Hong Wei, Yi Xiong, Jianyi Yang, Shanfeng Zhu

## Abstract

Homologous protein search is one of the most commonly used methods for protein annotation and analysis. Compared to structure search, detecting distant evolutionary relationships from sequences alone remains challenging. Here we propose PLMSearch (**P**rotein **L**anguage **M**odel), a homologous protein search method with only sequences as input. With deep representations from a pre-trained protein language model to predict similarity, PLMSearch can capture the remote homology information hidden behind the sequences. Extensive experimental results show that PLMSearch can search millions of query-target protein pairs in seconds like MMseqs2 while increasing the sensitivity by more than threefold, and is comparable to state-of-the-art structure search methods. In particular, unlike traditional sequence search methods, PLMSearch can recall most remote homology pairs with low sequence similarity but sharing similar structures. PLMSearch is freely available at https://dmiip.sjtu.edu.cn/PLMSearch.

## 1 Introduction

Homologous protein search is a key component of bioinformatics methods used in protein function prediction [1–5], protein-protein interaction prediction [6], and protein-phenotype association prediction [7]. The goal of homologous protein search is to find the homologous proteins from the target dataset (generally a large-scale standard dataset like Swiss-Prot [8]) for each query protein. The target protein with a higher probability of homology should be ranked higher. According to the type of input data, homologous protein search can be divided into sequence search and structure search.

Due to the low cost and large scale of sequence data, the most widely used homologous protein search methods are based on sequence similarity, such as MMseqs2 [9], BLASTp [10], and Diamond [11]. Despite the success of homology inference based on sequence similarity, it remains challenging to detect distant evolutionary relationships from sequences only [12]. Sequence profiles and profile hidden Markov models (HMMs) are condensed representations of multiple sequence alignment (MSAs) that specify for each position the probability of observing each of the 20 amino acids in evolutionarily related proteins. When the sequence identity is lower than 0.3, methods based on profile HMMs such as HMMER [13], HHsearch [14], and HHblits [15, 16] are currently the best tools for homologous protein search.

In cases of highly distant evolutionary relationships, sequences may have diverged to such an extent that we can no longer detect their relatedness. Since structures diverge much more slowly than sequences, detecting similarity between protein structures by 3D superposition provides higher sensitivity [17]. Protein structure search methods can be divided into: (1) contact/distance map-based, such as Map align [18], Eigen-THREADER [19], and DiscoVER [20]; (2) structural alphabet-based, such as 3D-BLAST-SW [21], CLE-SW [22], Foldseek, and Foldseek-TM [23]; (3) structural alignment-based, such as CE [24], Dali [25], and TM-align [26, 27]. Protein structure prediction methods (like AlphaFold2) and AlphaFold Protein Structure Database (AFDB) have greatly reduced the cost of obtaining protein structures [28–30], which expands the usage scenarios of the structure search methods. However, in the vast majority of cases, the sequence search method is still more convenient. This is notably evident in scenarios involving a large number of new sequences, such as metagenomic sequences [31], sequences generated by protein engineering [32], and antibody variant sequences [33].

At the same time, protein language models (PLMs) such as ESMs [34–36] and ProtTrans [37] only take protein sequences as input, trained on hundreds of millions of unlabeled protein sequences using self-supervised tasks such as masked amino acid prediction. PLMs perform well in various downstream tasks [38], especially in structure-related tasks like secondary structure prediction and contact prediction [39]. More recently, ProtENN [40] uses an ensemble deep learning framework that generated protein sequence embeddings to classify protein domains into Pfam families [41]; CATHe [42] trains an ANN on embeddings from the PLM ProtT5 [37] to detect remote homologs for CATH [43] superfamilies; Embedding-based annotation transfer (EAT) [44] uses Euclidean distance between vector representations (ProtT5 embeddings) of proteins to transfer annotations from a set of labeled lookup protein embeddings to query protein embeddings; DEDAL [45] and latest pLM-BLAST [46] obtain a continuous representation of protein sequences that, combined with the Smith-Waterman (SW) or Needleman-Wunsch (NW) algorithm, leads to a more accurate pairwise sequence alignment and homology detection method. These methods apply representations generated by deep learning models to protein domain classification, protein annotation, and pairwise sequence alignment, fully demonstrating the advantage of deep learning models in identifying remote homology. However, protein language models are not fully utilized for the large-scale protein sequence search.

To improve the sensitivity while maintaining the universality and efficiency of sequence search, we propose PLMSearch (Fig. 1a-c). PLMSearch mainly consists of the following three steps: (1) PfamClan filters out protein pairs that share the same Pfam clan domain [41]. (2) SS-predictor (**S**tructural **S**imilarity predictor) predicts the similarity between all query-target pairs with embeddings generated by the protein language model.

**Fig. 1.**
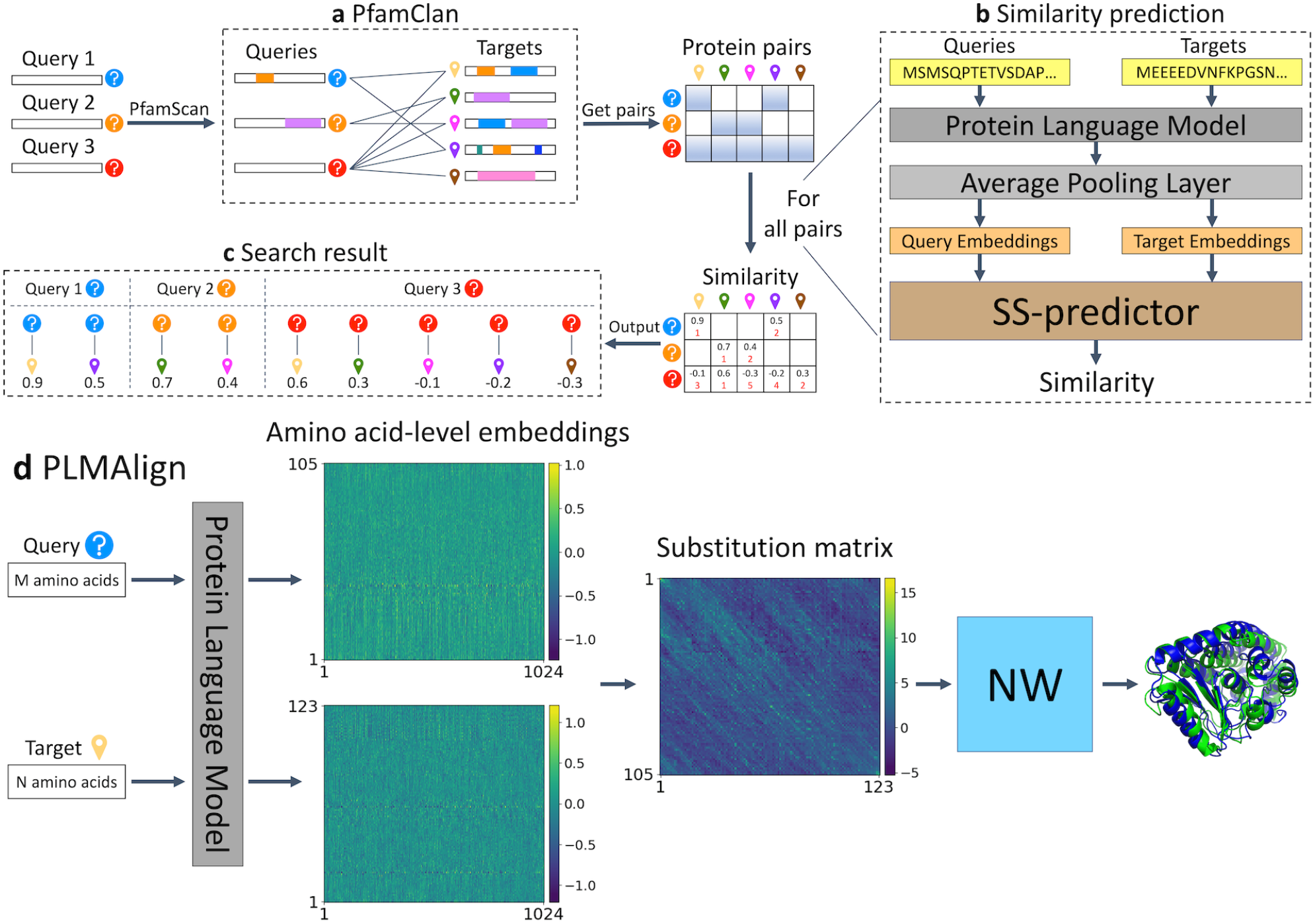
Overview of the PLMSearch pipeline. **a**, PfamClan. Initially, PfamScan [50] identifies the Pfam clan domains of the query protein sequences, which are depicted in different color blocks. Subsequently, PfamClan searches the target dataset for proteins sharing the same Pfam clan domain with the query proteins. Notably, the last query protein lacks any Pfam clan domain, and therefore, its all pairs with target proteins are retained. **b**, Similarity prediction. The protein language model generates deep sequence embeddings for query and target proteins. Subsequently, SS-predictor predicts the similarity of all query-target pairs. **c**, Search result. Finally, PLMSearch selects the similarity of the protein pairs prefiltered by PfamClan, sorts these protein pairs based on their predicted similarity, and outputs the search results for each query protein separately. **d**, PLMAlign. PLMAlign utilizes per-residue embeddings as input to compute a substitution matrix. This substitution matrix is then employed to replace the static substitution matrix in the Smith-Waterman (SW) or Needleman-Wunsch (NW) algorithm, enabling the local or global sequence alignment. The global alignment is illustrated in the figure, where the length of the query protein is 105, the length of the target protein is 123, and the embedding dimension of ProtT5-XL-UniRef50 used by PLMAlign is 1024.

PLMSearch will not lose much sensitivity without structures as input, because it uses the protein language model to capture remote homology information from deep sequence embeddings. In addition, the SS-predictor used in this step uses the structural similarity (TM-score) as the ground truth for training. This allows PLMSearch to acquire reliable similarity even without structures as input. (3) PLMSearch sorts the pairs pre-filtered by PfamClan based on their predicted similarity and outputs the search results for each query protein accordingly. redSubsequently, PLMAlign provides sequence alignments and alignment scores for top-ranked protein pairs retrieved by PLMSearch (Fig. 1d). Search tests on SCOPe40-test and Swiss-Prot reveal that PLM-Search is always one of the best methods and provides the best tradeoff between accuracy and speed. Specifically, PLMSearch can search millions of query-target protein pairs in seconds like MMseqs2, but increases the sensitivity by more than threefold, and approaches the state-of-the-art structure search methods. The improvement in sensitivity is particularly apparent in remote homology pairs.

## 2 Results

### 2.1 PLMsearch reaches similar sensitivities as structure search methods

We benchmarked the sensitivity of SS-predictor, PLMSearch, PLMSearch + PLMAlign, five other sequence search methods (MMseqs2, Blastp, HHblits, EAT, and pLM-BLAST), four structural alphabet-based search methods (3D-BLAST-SW, CLE-SW, Foldseek, and Foldseek-TM), and three structural alignment-based search methods (CE, Dali, and TM-align). We performed search tests on SCOPe40-test and Swiss-Prot after filtering homologs from the training dataset (see “Datasets”, “Metrics”, and “Baselines” Section). In the SCOPe40-test dataset (2,207 proteins), we performed an all-versus-all search test. Therefore, a total of 4,870,849 query-target pairs were tested for all the methods. Fig. 2a-c shows the results of the 11 most competitive methods in sensitivity and speed. Supplementary Fig. 1 shows the results of the other two structural alphabet-based and two structural alignment-based search methods. In the Swiss-Prot search test, we randomly selected 50 queries from Swiss-Prot and 50 queries from SCOPe40-test (a total of 100 query proteins) and searched for 430,140 target proteins in Swiss-Prot. Therefore, a total of 43,014,000 query-target pairs were tested for the six most efficient methods on searching large-scale datasets (Supplementary Fig. 2, Supplementary Table 2).

**Fig. 2.**
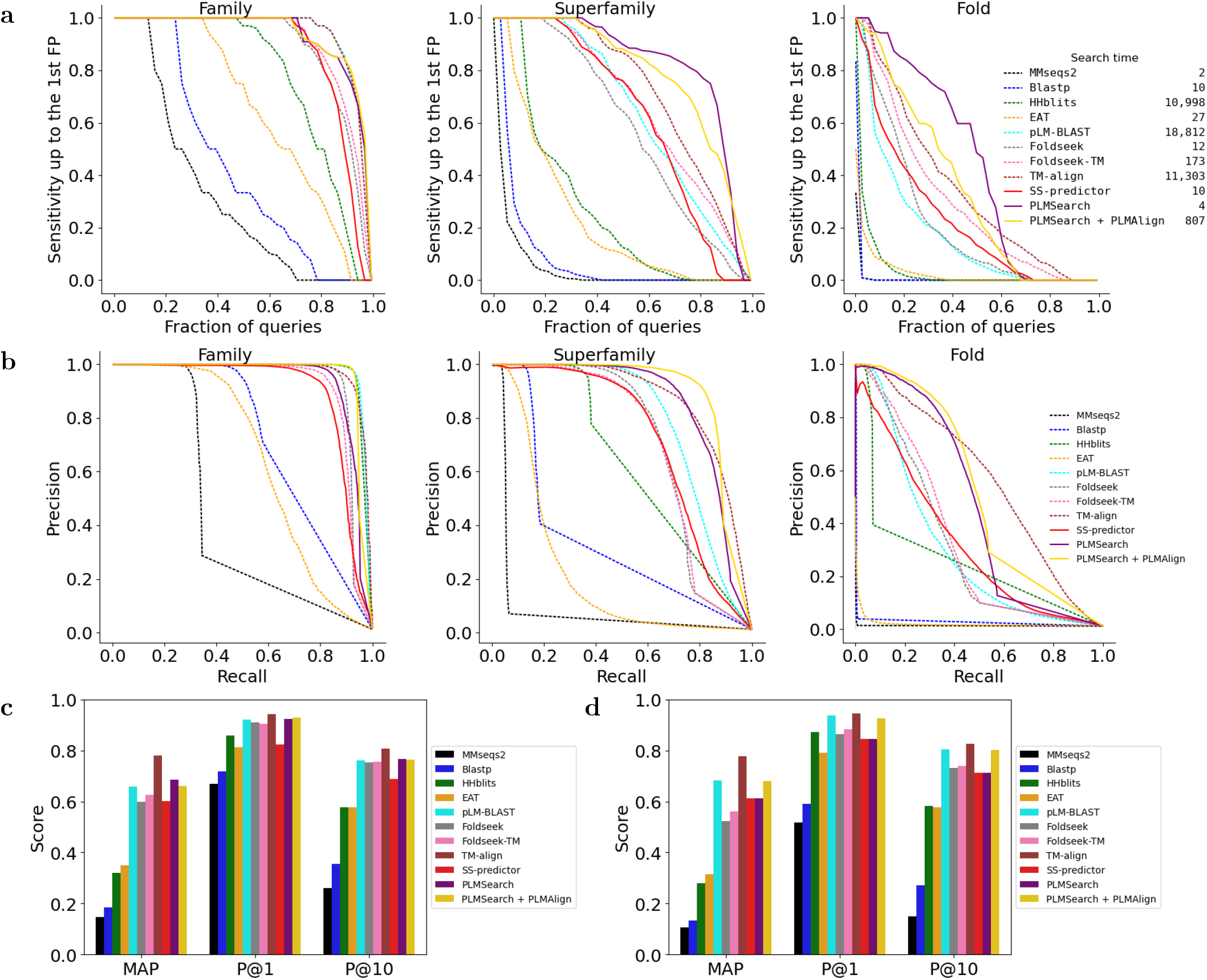
PLMsearch reaches similar sensitivities as structure search methods. **a-c**, The all-versus-all search test on SCOPe40-test. For family-level, superfamily-level, and fold-level recognition, TPs were defined as same family, same superfamily but different family, and same fold but different superfamily, respectively. Hits from different folds are FPs. After sorting the alignment result of each query according to similarity, we calculated the sensitivity as the fraction of TPs in the sorted list up to the first FP to better reflect the requirements for low false discovery rates in automatic searches. **a**, We took the mean sensitivity over all queries as AUROC. In addition, the total search time for the all-versus-all search test with a 56-core Intel(R) Xeon(R) CPU E5-2680 v4 @ 2.40 GHz and 256 GB RAM server is shown on the legend. **b**, Precision-recall curve. **c**, MAP, P@1, and P@10. **d**, Evaluation on new proteins (see “New protein search test” Section). Extended Data Table A1 and Extended Data Table A3 record the specific values of each metric.

Extended Data Table A1 records the specific values of each metric. PLMSearch performs well on most metrics, especially at the superfamily-level and fold-level, which are shallower and have less significant similarity between protein sequences. PLMSearch is 3, 16, and 219 times exceeding MMseqs2 in AUROC of the family-level (from 0.318 to 0.928), superfamily-level (from 0.050 to 0.826), and fold-level (from 0.002 to 0.438), respectively. Extended Data Table A2 indicates that the primary reason for the improvement is that PLMSearch is more robust and makes the rank of the first FP (**F**alse **P**ositive) lower, increasing the number of total TPs (**T**rue **P**ositives) up to the first FP (38 times exceeding MMseqs2, from 2.74 to 104.78). PLMSearch + PLMAlign uses PLMAlign to align protein pairs with a similarity exceeding 0.3 from PLMSearch. The alignment score is then used to rerank, which improves the precision rate, especially on the search test for new proteins that fail to scan out any Pfam domain (Fig. 2d, Extended Data Table A3). Accordingly, the exclusion of pairs with a similarity below 0.3 leads to a marginal decrease in the recall rate.

### 2.2 PLMSearch searches millions of query-target pairs in seconds

We first compared the total search time of different methods for the all-versus-all search test on SCOPe40-test (2,207 proteins, 4,870,849 query-target pairs). To ensure fairness, we used the same computing resources (a 56-core Intel(R) Xeon(R) CPU E5-2680 v4 @ 2.40 GHz and 256 GB RAM server) when implementing different methods in the evaluation. For HHblits, pLM-BLAST, TM-align, and various other parallelizable methods, we utilized all 56 cores by default. Moreover, almost all methods need to preprocess the target dataset in advance. This part of the time is not required while searching, so we did not include the preprocessing time in the search time statistics. As shown in the legend of Fig. 2a, by using SS-predictor to predict the similarity instead of calculating the structural similarity (TM-score) of all protein pairs, SS-predictor (10 s) and PLMSearch (4 s) are one of the fastest methods, and more than four orders of magnitude faster than TM-align (11,303 s).

PLMSearch can achieve similar efficiency on our publicly available web server with CPU only (64 * Intel(R) Xeon(R) CPU E5-2682 v4 @ 2.50 GHz and 512 GB RAM). Searching a query against Swiss-Prot (568K proteins) and UniRef50 (53.6M proteins) and employing PLMAlign to align the query with the Top-10 targets [8] requires approximately 0.15 minutes and 1.1 minutes, respectively (Extended Data Table A4). In fact, when searching a query against Swiss-Prot (568K proteins), PLMAlign takes up 0.12 minutes (more than 80% of the total time), and PLMSearch only takes about 0.03 minutes (Supplementary Table 1). This is because PLMSearch generates and preloads the embeddings of all target proteins in advance. This strategy helps to save much time by avoiding repeated forward propagations of the protein language model with a large number of parameters and saves the time for loading embeddings from disk to RAM. As a result, just one forward pass through the SS-predictor network is required to predict the similarity of millions of query-target pairs and complete the search in seconds (Fig. 5b).

### 2.3 PLMSearch accurately detects remote homology pairs

Remote homology pairs generally refer to homologous protein pairs with dissimilar sequences but similar structures [47]. Such protein pairs have low sequence similarity, so their homology is difficult to be detected by sequence alignment-based methods (MMseqs2, Blastp), but can be detected by structure-based search methods (Foldseek, Foldseek-TM) (Fig. 3a). In this study, pairs with similar sequences and similar structures are defined as sequence identity > 0.3 [47] and TM-score > 0.5 [48, 49] and are called “easy pairs”; pairs with dissimilar sequences but similar structures are defined as sequence identity < 0.3 but TM-score > 0.5 and are called “remote homology pairs”. We conducted a specific analysis of recalled pairs and missed pairs (defined in Fig. 3b). We calculated the TM-score and the sequence identity (see “Sequence alignment” Supplementary Section) of the recalled/missed pairs and projected them onto a 2D scatter plot (Fig. 3c-h).

**Fig. 3.**
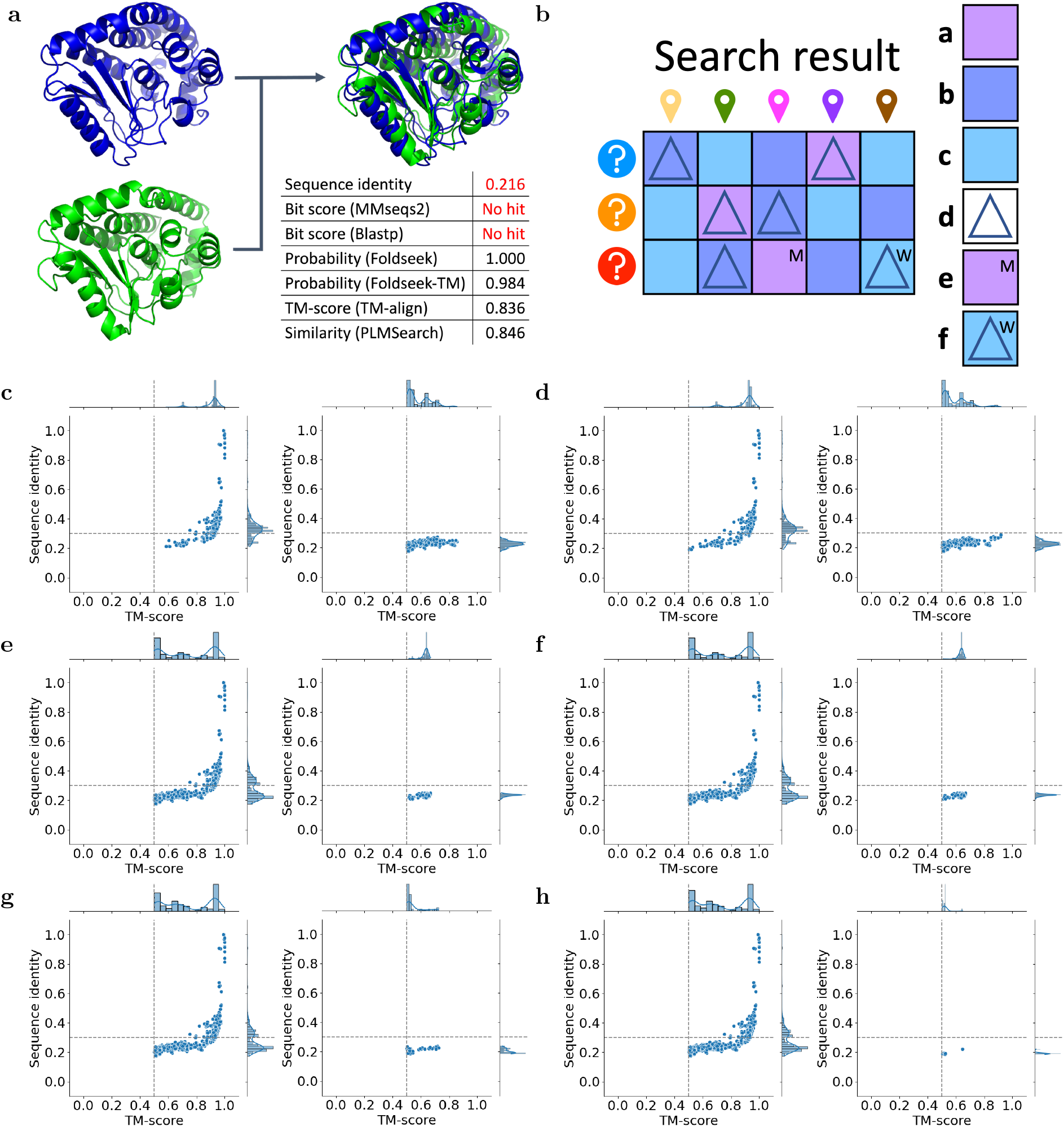
PLMSearch accurately detects remote homology pairs. **a**, Case study. Protein pairs with dissimilar sequences but similar structures are called remote homology pairs here. The sequence identity between the Q08558 (the blue structure) and I6Y3U6 (the green structure) is low (0.216<0.3). Thus, it is difficult to find this remote homology pair with sequence alignment only. However, Foldseek, Foldseek-TM, TM-align, and PLMSearch capture the remote homology pair that is missed by MMseqs2 and Blastp. **b**, Definition diagram. This search result example is composed of three queries and five targets. Among the 15 protein pairs, **Legend a** marks three pairs with a TM-score > 0.5, usually assumed to have the same fold [48, 49]. **Legend b** marks six pairs with a TM-score between 0.2 and 0.5. **Legend c** marks six pairs with a TM-score < 0.2, usually assumed as randomly selected irrelevant pairs [48, 49]. **Legend d** marks six filtered pairs. **Legend e** marks the pair at (3,3) with a TM-score > 0.5 but is not filtered out, which is a “Missed” pair. Correspondingly, protein pairs in (1,4) and (2,2) are “Recalled” pairs. **Legend f** marks the pair at (3,5) with a TM-score < 0.2 but is filtered out, which is a “Wrong” pair. **c-h**, From the search results of five randomly selected queries to avoid oversampling (with Swiss-Prot as the target dataset, a total of 2,150,700 query-target pairs), we selected the 5000 pairs with the highest similarity for different search methods and counted the recalled and missed pairs: **c**, MMseqs2. **d**, Blastp. **e**, Foldseek. **f**, Foldseek-TM. **g**, SS-predictor. **h**, PLMSearch. For recalled pairs (left) and missed pairs (right) in each subplot, the TM-score (x-axis) and sequence identity (y-axis) between protein pairs are shown on the 2D scatter plot. The thresholds, sequence identity = 0.3 [47] and TM-score = 0.5 [48, 49], are shown by dashed lines. All methods successfully recall the easy pairs in the first quadrant. But for remote homology pairs in the fourth quadrant, SS-predictor & PLMSearch did the best, followed by Foldseek & Foldseek-TM, and MMseqs2 & Blastp were the worst. Extended Data Table A5 records the specific values of each metric.

Compared with easy pairs (the first quadrant in Fig. 3c-h), remote homology pairs (the fourth quadrant in Fig. 3c-h) in the “twilight zone” of protein sequence homology are more difficult to detect [47]. Among the six methods (Extended Data Table A5), even the least sensitive methods MMseqs2 and Blastp recall all the easy pairs (574/574), but perform poorly on remote homology pairs (MMseqs2: 183/1105, Blastp: 203/1105). In contrast, powered by the protein language model, SS-predictor and PLM-Search search out most of the remote homology pairs (SS-predictor: 1022/1105, PLMSearch: 1087/1105, six times exceeding MMseqs2), and the recall rate exceeds Foldseek, which directly uses structural data as input (Foldseek: 934/1105, Foldseek-TM: 940/1105).

### 2.4 Ablation experiments: PfamClan, SS-predictor, and PLMAlign make PLMSearch more robust

To evaluate PLMSearch without the PfamClan component, we screened a total of 110 queries from the 2,207 queries in SCOPe40-test, which failed to scan any Pfam domain (see “New protein search test” Section). As expected, the performance of PLMSearch is exactly the same as that of SS-predictor, because PfamClan does not filter out any protein pairs, whereas PLMSearch still produces relatively sensitive search results (MAP = 0.612, P@1 = 0.845, P@10 = 0.712, see Extended Data Fig. A1e, Extended Data Table A3). Using PLMAlign to align and rank based on alignment scores significantly enhances precision. This improvement stems from the fact that, unlike SS-predictor, PLMAlign employs per-residue embeddings rather than per-protein embeddings as input and uses pairwise alignment instead of large-scale similarity prediction. Besides, it is noteworthy that both SS-predictor + PLMAlign and PLM-Search + PLMAlign only align pairs from SS-predictor and PLMSearch pre-filter results with a similarity exceeding 0.3 (totaling 1,591,492 and 379,707 pairs, respectively), in contrast to aligning all pairs like PLMAlign/pLM-BLAST (4,870,849). This streamlined approach significantly reduces the alignment time (nearly 16 times) while maintaining comparable precision, underscoring the benefits of leveraging SS-predictor and PLM-Search to pre-filter (Extended Data Fig. A1b).

To evaluate PLMSearch without the SS-predictor component, we first clustered the SCOPe40-test and Swiss-Prot datasets based on PfamClan (Extended Data Fig. A2, Supplementary Table 3). Specifically, proteins belonging to the same Pfam clan are clustered. The clustering results show a significant long-tailed distribution. After pre-filtering with PfamClan, more than 50% of the pre-filtered protein pairs (orange rectangles in the figure) are from the largest 1-2 clusters (big clusters), which only accounts for a very small part of the entire clusters (SCOPe40-test: 0.231%; Swiss-Prot: 0.032%). Therefore, big clusters will result in a significant number of irrelevant protein pairs in the pre-filtering results, reducing accuracy, and must be further sorted and filtered based on similarity, which is what SS-predictor does.

Furthermore, among all similarity-based search methods (See “Baselines”), we further compared the correlation between the predicted similarity and TM-score (Extended Data Fig. A1a). The correlation between the similarity predicted by Euclidean (COS) and TM-score is not high, resulting in a large number of actually dissimilar protein pairs ranking first. The similarity predicted by SS-predictor is more correlated with TM-score (with a higher Spearman correlation coefficient). This is why SS-predictor outperforms other similarity-based search methods with the same embeddings as input (Extended Data Fig. A1b-e, Extended Data Table A1, Extended Data Table A3).

## 3 Discussion

We study the use of protein language models for large-scale homologous protein search in this work. We propose PLMSearch, which takes only sequences as input and searches for homologous proteins using the protein language model and Pfam sequence analysis, allowing PLM-Search to extract remote homology information buried behind sequences. Subsequently, PLMA-lign is used to align protein pairs retrieved by PLMSearch and obtain the alignment scores. Experiments reveal that PLMSearch outperforms MMseqs2 in terms of sensitivity and is comparable to the state-of-the-art structural search approaches. The improvement is especially noticeable in remote homology pairs. PLMSearch, on the other hand, is one of the fastest search methods in comparison to other baselines, capable of searching millions of query-target protein pairs in seconds. We also summarized the 11 most competitive approaches based on their input and performance in Supplementary Table 4.

We discuss the differences between search methods (like PLMSearch) and alignment methods (like pLM-BLAST and PLMAlign) in detail in Supplementary Table 5. It is noteworthy that residue embedding-based alignment methods, such as PLMAlign and pLM-BLAST [46], achieve respectable sensitivity. However, a primary limitation lies in the maximum size of the target dataset. This is particularly evident in two key aspects: (1) Residue embedding-based alignment necessitates retaining the embeddings of all residues for each protein in the target dataset, denoted as *N* \**L*_*i*_ \**D* (where *N* is the number of proteins, *L*_*i*_ is the length of the protein, and *D* is the embedding dimension). In contrast, PLMSearch only requires retaining per-protein embeddings, expressed as *N* * 1 * *D*. This results in a size difference exceeding three orders of magnitude, posing a significant challenge when implementing a dataset with the size of UniRef50, which contains 53.6 million proteins [8]. (2) Residue embedding-based alignment determines the similarity between protein pairs through pairwise global (local) alignments. In contrast, PLMSearch only needs a single forward pass through the SS-predictor network to predict the similarity of millions of query-target pairs. However, it is important to note that PLMSearch can solely predict the similarity of protein pairs without any alignment suggestions. To this end, PLMSearch + PLMAlign provides alignment for protein pairs filtered by PLMSearch with a similarity higher than 0.3. This approach not only compensates for PLMSearch’s limitations but also avoids numerous low similarity and meaningless alignments, thereby maintaining high efficiency. In the future, we intend to explore the mutual attention between query and target per-residue embeddings to provide better global and local sequence alignment results.

In summary, we believe that PLMSearch has removed the low sensitivity limitations of sequence search methods. Since the sequence is more applicable and easier to obtain than structure, PLM-Search is expected to become a more convenient large-scale homologous protein search method.

## 4 Methods

### 4.1 PLMSearch pipeline

PLMSearch consists of three steps (Fig. 1a-c). (1) PfamClan. Initially, PfamScan [50] identifies the Pfam clan domains of the query protein sequences. Subsequently, PfamClan searches the target dataset for proteins sharing the same Pfam clan domain with the query proteins. In addition, a limited number of query proteins lack any Pfam clan domain, or their Pfam clans differ from any target protein. To prevent such queries from yielding no results, all pairs between such query proteins and target proteins will be retained. (2) Similarity prediction. The protein language model generates deep sequence embeddings for query and target proteins. Subsequently, SS-predictor predicts the similarity of all query-target pairs. (3) Search result. Finally, PLMSearch selects the similarity of the protein pairs prefiltered by PfamClan, sorts the protein pairs based on their similarity, and outputs the search results for each query protein separately.

For the top-ranked query-target pairs, PLMA-lign is used to generate local or global alignments and alignment scores. In addition, we also added parallel sequence-based Needleman-Wunsch alignment and structure-based TM-align at the end of our pipeline for users to choose.

### 4.2 PfamClan

PfamClan filters out protein pairs that share the same Pfam clan domain (Fig. 1a). It is worth noting that the recall rate is more important in the initial pre-filtering. PfamClan is based on a more relaxed standard of sharing the same Pfam clan domain, instead of sharing the same Pfam family domain (what PfamFamily does). This feature allows PfamClan to outperform PfamFamily in recall rate (Fig. 4) and successfully recalls high TM-score protein pairs that PfamFamily misses (Extended Data Table A6).

**Fig. 4.**
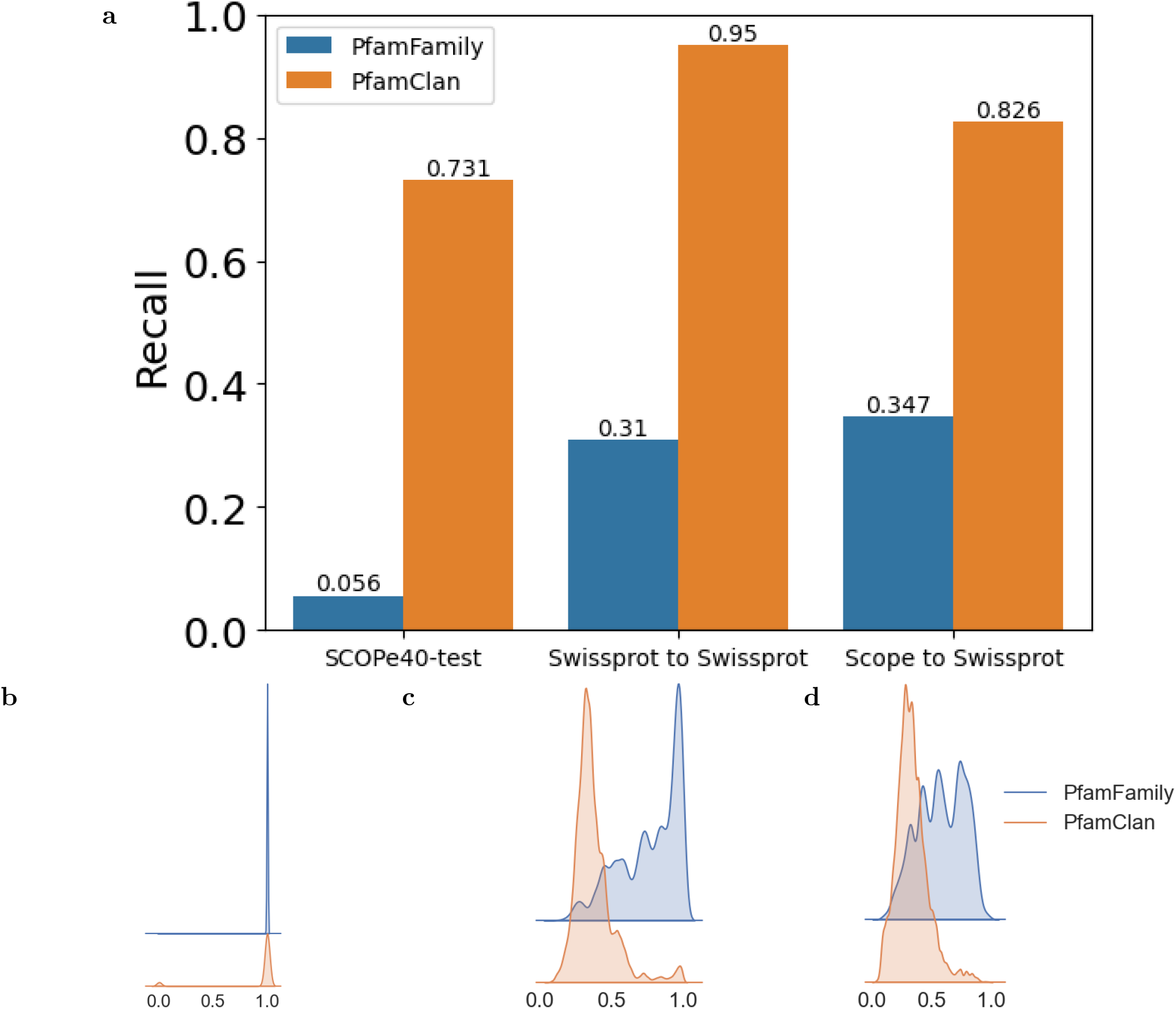
The pre-filtering results of PfamFamily & PfamClan. **a**, We evaluated the pre-filtering results of PfamFamily & PfamClan on the SCOPe40-test, Swiss-Prot to Swiss-Prot, and SCOPe40-test to Swiss-Prot search tests (see “Datasets”). PfamClan achieves a higher recall rate. **b**, Same(1) or Different(0) fold on SCOPe40-test, **c-d**, TM-score distributions using kernel density estimation (smoothed histogram using a Gaussian kernel with the width automatically determined). **c**, Swiss-Prot to Swiss-Prot; **d**, SCOPe40-test to Swiss-Prot. The distribution of PfamFamily is to the right as a whole, because the requirements of PfamFamily are stricter than PfamClan, so the protein pair it recalls has a higher probability of being in the same fold and sharing a higher TM-score. However, this also leads to PfamFamily having a lower recall rate and missing some homologous protein pairs as shown in Extended Data Table A6. It is worth noting that the recall rate is more important in the initial pre-filtering.

### 4.3 Similarity prediction

Based on the protein language model and SS-predictor, PLMSearch performs further similarity prediction based on the pre-filtering results of PfamClan (Fig. 1b). The motivation is that the clustering results based on PfamClan show a significant long-tailed distribution (Extended Data Fig. A2). As the size of the dataset increases, the number of proteins contained in big clusters will greatly expand, further leading to a rapid increase in the number of pre-filtered protein pairs (Supplementary Table 3). The required computing resources are excessive with TM-align used for all the filtered pairs. PLMSearch uses the predicted similarity instead, which helps to greatly increase speed and avoids dependence on structures.

As shown in Fig. 5a, the input protein sequences are first sent to the protein language model (ESM-1b here) to generate per-residue embeddings (*m* * *d*, where *m* is the sequence length and *d* is the dimension of the vector), and the per-protein embedding (1 * *d*) is obtained through the average pooling layer. Subsequently, SS-predictor predicts the structural similarity (TM-score) between proteins through a bilinear projection network (Fig. 5b). However, it is difficult for the predicted TM-score to directly rank protein pairs with extremely high sequence identity (often differing by only a few residues). This is because the predicted TM-score was trained on SCOPe40-train and CATHS40, where protein pairs share sequence identity < 0.4, so cases with extremely high sequence identity were not included. At the same time, COS similarity performs well in cases with extremely high sequence identity (see P@1 in Extended Data Fig. A1d-e) but becomes increasingly insensitive to targets after Top-10. Therefore, the final similarity predicted by SS-predictor is composed of the predicted TM-scores and the COS similarity to complement each other. We studied the reference similarity of COS similarity in SCOPe40-train (Supplementary Fig. 3a-b, Supplementary Table 6). We assemble the predicted TM-score with the top COS similarity as follows. When COS similarity > 0.995, SS-predictor similarity equals COS similarity. Otherwise, SS-predictor similarity equals the predicted TM-score multiplied by COS similarity.

**Fig. 5.**
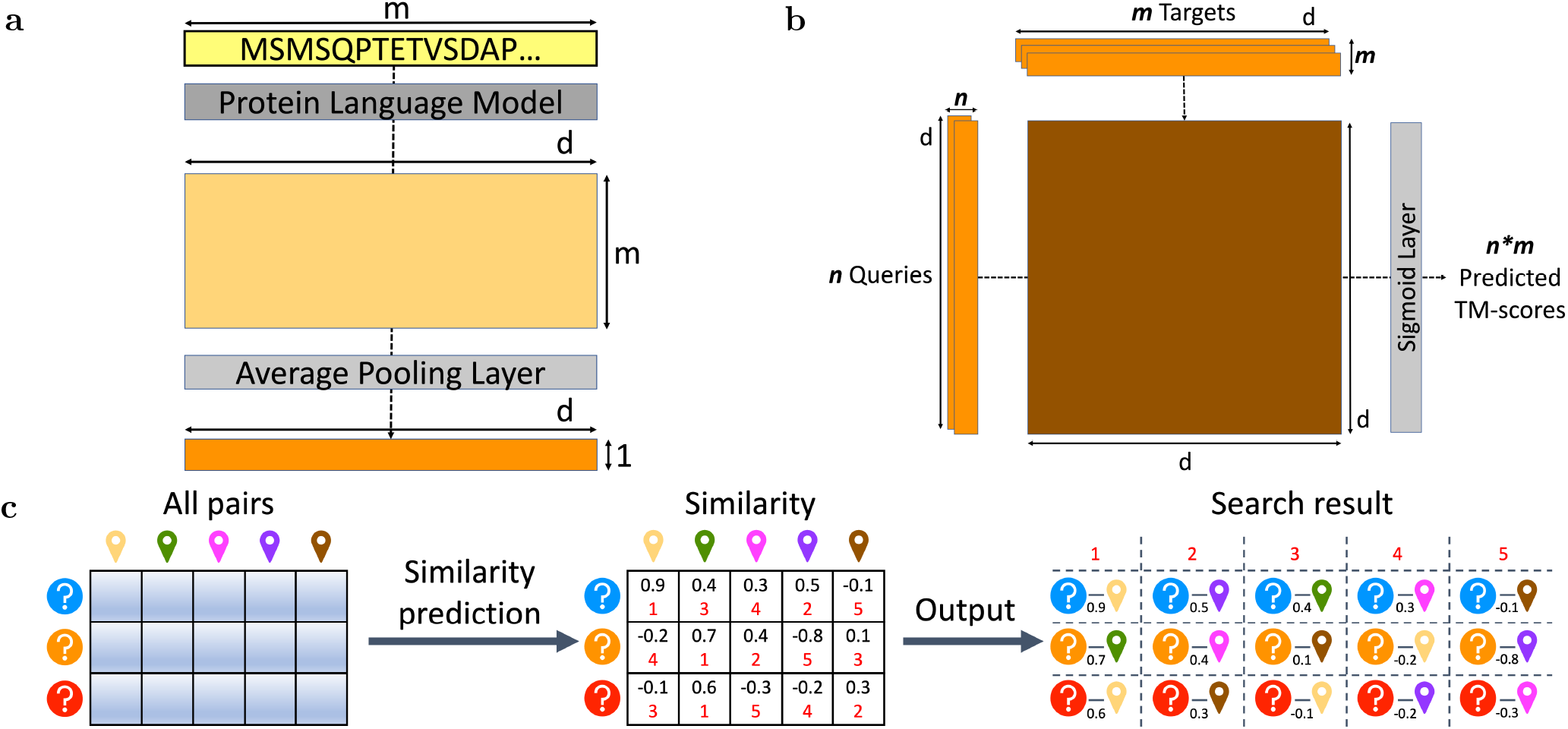
SS-predictor. **a**, Protein embedding generation. The protein language model converts the protein sequence composed of *m* residues into *m* * *d* per-residue embeddings, where *d* is the dimension of the embedding. Then the average pooling layer converts them into the corresponding 1 * *d* per-protein embedding *z*. **b**, SS-predictor. A bilinear projection network predicts all the TM-scores between *n* query embeddings *z*1 and *m* target embeddings *z*2 by *z*1 * *W* * *z*2, where the matrices *W* are the learned parameters. **c**, Similarity-based search methods. The similarity of all protein pairs is predicted and sorted and then outputted as a search result. According to different similarity, three corresponding search methods are: Euclidean, COS, and SS-predictor.

We also studied the reference similarity of SS-predictor similarity (Fig. 6) in SCOPe40-train. There is a clear phase transition occurring around the similarity of 0.3 − 0.7. Supplementary Table 6 shows that for SS-predictor, protein pairs with a similarity lower than 0.3 are usually assumed as randomly selected irrelevant protein pairs. In the ridge plot Fig. 6b, as expected, the protein pairs in the same fold and different folds are well grouped in two different similarity ranges, i.e. the protein pairs in the same fold have a higher similarity and the protein pairs in different folds have a lower one. However, since similarity and SCOP fold do not have a one-to-one correspondence, there is a small overlap. Furthermore, unlike Foldseek, which focuses on local similarity, the similarity of SS-predictor, like TM-score, focuses on global similarity (Extended Data Table A7).

**Fig. 6.**
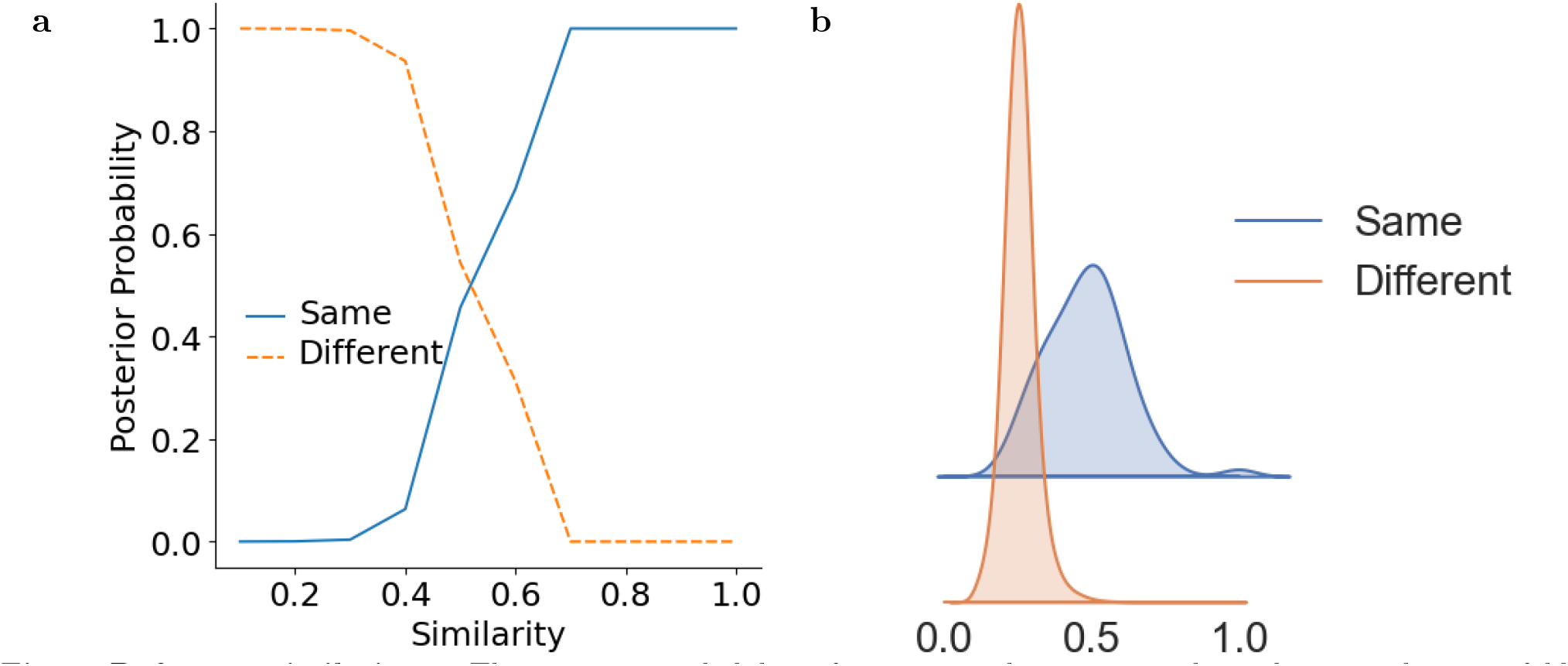
Reference similarity. **a**, The posterior probability of proteins with a given similarity being in the same fold or different folds in SCOPe40-train. **b**, Similarity distribution of the same and different folds protein pairs using kernel density estimation (smoothed histogram using a Gaussian kernel with the width automatically determined). The similarity of protein pairs belonging to the same protein pair is significantly higher than that of protein pairs belonging to different folds. The posterior probability corresponding to the similarity is shown in Supplementary Table 6. Protein pairs with a similarity lower than 0.3 is usually assumed as randomly selected irrelevant protein pair. See “Reference similarity” Supplement Section for more details.

### 4.4 PLMAlign

For the retrieved protein pairs, PLMAlign takes per-residue embeddings as input to obtain specific alignments and alignment scores (Fig. 1d). PLMAlign subsequently uses alignment scores to rerank, which improves the ranking results further. Specifically, inspired by pLM-BLAST [46], PLMAlign calculates the substitution matrix by dot producting the per-residue embeddings of the query-target protein pair. The substitution matrix is then used in the SW/NW algorithm to perform local/global alignment, and the algorithm is accelerated through the linear gap penalty. Compared with traditional SW/NW using a fixed substitution matrix, the substitution matrix calculated by PLMAlign using protein embedding generated from the sequence context, thus containing deep evolutionary information. Compared with pLM-BLAST, by using the dot product and the linear gap penalty, PLMAlign can better align remote homology pairs while reducing the algorithm complexity to *O*(*mn*) to ensure high efficiency (Supplementary Table 7). Therefore, PLMAlign performs better on remote homology alignment (using “Malisam and Malidup” datasets as benchmarks, see Supplementary Fig. 6, Supplementary Table 8). Also see “Remote homology alignment” Supplementary Section for detailed settings for PLMAlign and the evaluation of alignment results. The reference score of PLMAlign is provided in Supplementary Fig. 3c-d and Supplementary Table 6. PLMAlign in the main text uses global alignment to generate alignment scores.

### 4.5 Datasets

The data volumes and uses of each dataset are summarized in Extended Data Table A8.

#### 4.5.1 SCOPe40

The SCOPe40 dataset consists of single domains with real structures. Clustering of SCOPe

2.01 [51, 52] at 0.4 sequence identity yielded 11,211 non-redundant protein domain structures (“SCOPe40”). As done in Foldseek, domains from SCOPe40 were split 8:2 by fold into SCOPe40-train and SCOPe40-test, and then domains with a single chain were reserved. It is worth mentioning that each domain in SCOPe40-test belongs to a different fold from all domains in SCOPe40-train, so the difference between training and testing data is much larger than that of pure random division. We also studied the max sequence identity of each protein in SCOPe40-test relative to the training dataset (Supplementary Fig. 4). From the figure, we can draw a similar conclusion that the sequences in SCOPe40-test and the training dataset are quite different, and most of the max sequence identity is between 0.2 and 0.3. PLM-Search performs well in SCOPe40-test (Fig. 2, Extended Data Table A1), implying that PLM-Search learns universal biological properties that are not easily captured by other sequence search methods [12].

#### 4.5.2 New protein search test

In real-world scenarios, newly discovered and unclassified proteins often play a crucial role in innovative research. To assess the effectiveness of various methods in searching these proteins, we introduced an additional search test exclusively comprising query proteins that failed to scan any Pfam domain. Specifically, we screened a total of 110 queries from the 2,207 queries in the SCOPe40-test, which failed to scan any Pfam domain. In the all-versus-all search test on the SCOPe40-test dataset, we counted the MAP, P@1, and P@10 metrics with only new proteins as queries (110 proteins) and SCOPe40-test as targets (2,207 proteins).

#### 4.5.3 Swiss-Prot

Unlike SCOPe, the Swiss-Prot dataset consists of full-length, multi-domain proteins with predicted structures, which are closer to real-world scenarios. Because the throughput of experimentally observed structures is very low and requires a lot of human and financial resources, the number of real structures in datasets like PDB [53–55] tends to be low. AlphaFold protein structure database (AFDB) obtains protein structure through deep learning prediction, so it contains the entire protein universe and gradually becomes the mainstream protein structure dataset. Therefore, in this set of tests, we used Swiss-Prot with predicted structures from AFDB as the target dataset.

Specifically, we downloaded the protein sequence from UniProt [8] and the predicted structure from the AlphaFold Protein Structure Database [30]. A total of 542,317 proteins with both sequences and predicted structures were obtained. For these proteins, we dropped low-quality proteins with an avg. pLDDT lower than 70, and left 498,659 proteins. In order to avoid possible data leakage issues, like SCOPe40, we used 0.4 sequence identity as the threshold to filter homologs in Swiss-Prot from the training dataset. Specifically, we used the previously screened 498,659 proteins as query proteins and SCOPe40-train as the target dataset. We first pre-filtered potential homologous protein pairs with MMseqs2 and calculated the sequence identity between all these pairs. The query protein from Swiss-Prot will be discarded if the sequence identity between the query protein and any target protein is greater than or equal to 0.4. Finally, 68,519 proteins were deleted via homology filtering, leaving 430,140 proteins in Swiss-Prot, which we employed in our experiments. We also studied the max sequence identity of each protein in Swiss-Prot relative to training dataset (Supplementary Fig. 4) and found that the vast majority of them were below 0.3.

Subsequently, we randomly selected 50 queries from Swiss-Prot and 50 queries from SCOPe40-test as query proteins (a total of 100 query proteins) and searched for 430,140 target proteins in Swiss-Prot. Therefore, a total of 43,014,000 query-target pairs were tested. The search test for 50 query proteins from Swiss-Prot and SCOPe40-test are called “Swiss-Prot to Swiss-Prot” and “SCOPe40-test to Swiss-Prot”, respectively.

#### 4.5.4 CATHS40

The SCOPe40-train dataset includes 8,953 proteins and TM-scores for all protein pairs were calculated for training. As the majority of these pairs had TM-scores below 0.5, only 504,553 pairs (among 80,156,209 in total) had a TM-score above 0.5 for model training. To enhance the model’s generality, we supplemented it with high-quality protein pairs extracted from the curated CATH domain dataset [43, 56]. We began with the CATHS40 non-redundant dataset of protein domains, which exhibits no more than 0.4 sequence similarity. Domains exceeding 300 residues were filtered out, leaving 27,270 domains. To prevent potential data leakage issues, akin to SCOPe40, we applied a 0.4 sequence identity threshold to filter homologs in CATHS40 from the testing dataset (SCOPe40-test and Swiss-Prot).

Finally, 21,474 proteins in CATHS40 were left for training, and the max sequence identity of the test dataset to the new training dataset is still less than 0.4 (Supplementary Fig. 4). We then undersampled CATHS40 domain pairs from different folds to acquire a substantial amount of training pairs with TM-scores above 0.5. Specifically, we sampled the TM-scores of 28,440,312 protein pairs for training, of which 7,813,946 pairs had a TM-score above 0.5 (Extended Data Table A8).

#### 4.5.5 Target datasets on web server

We currently have the following four target datasets for users to search: (1) Swiss-Port (568K proteins) [8], the original dataset without filtering; (2) PDB (680K proteins) [53–55]; (3) UniRef50 (53.6M proteins) [8]. UniRef50 is built by clustering UniRef90 seed sequences that have at least 50% sequence identity to and 80% overlap with the longest sequence in the cluster; (4) Self (the query dataset itself).

#### 4.5.6 Malisam and Malidup

To establish a robust reference for alignment, we used two gold-standard benchmark datasets known as Malisam [57] and Malidup [58]. These sets meticulously curate structural alignments through manual curation, emphasizing challenging-to-detect, low-sequence-identity remote homology. Malidup encompasses 241 pairwise structure alignments, specifically targeting homologous domains within the same chain, thereby exemplifying structurally similar remote homologs. Malisam consists of analogous motifs.

### 4.6 Metrics

We evaluated different homologous protein search methods using the following four metrics: AUROC, AUPR, MAP, and P@K.

- AUROC [23]: The mean sensitivity over all queries, where the sensitivity is the fraction of TPs in the sorted list up to the first FP, all excluding self-hits.
- AUPR [23]: Area under the precision-recall curve.
- MAP [59]: Mean average precision (MAP) for a set of queries is the mean of the average precision scores for all query proteins.

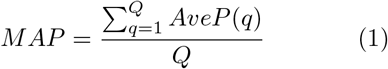

where Q is the number of queries.
- P@K [19]: For homologous protein search, as many queries have thousands of relevant targets, and few users will be interested in getting all of them. Precision at k (P@k) is then a useful metric (e.g., P@10 corresponds to the number of relevant results among the top 10 retrieved targets). P@K here is the mean value for each query.

On SCOPe40-test, we performed an all-versus-all search test, which means both the query and the target dataset were SCOPe40-test. To make a more objective comparison, the settings used in the all-versus-all search test are exactly the same as those used in Foldseek [23]. Specifically, for family-level, superfamily-level, and fold-level recognition (Supplementary Fig. 5), TPs were defined as the same family, same superfamily but different family, and same fold but different super-family, respectively. Hits from different folds are FPs. After sorting the search result of each query according to similarity (described in Supplementary Table 9), we calculated the sensitivity as the fraction of TPs in the sorted list up to the first FP to better reflect the requirements for low false discovery rates in automatic searches. We then took the mean sensitivity over all queries as AUROC. Additionally, we plotted weighted precision-recall curves (precision = TP/(TP+FP) and recall = TP/(TP+FN)). All counts (TP, FP and FN) were weighted by the reciprocal of their family, superfamily or fold size. In this way, families, superfamilies and folds contribute linearly with their size instead of quadratically [13]. MAP and P@K were calculated according to the TPs and FPs defined by fold-level.

On search tests against Swiss-Prot, TPs were defined as pairs with a TM-score > 0.5, otherwise FPs. MAP and P@K are then calculated accordingly. The reason is that it is problematic to evaluate the SCOPe domain annotations of multidomain proteins on search tests against Swiss-Prot [23]. Without the Astral SCOPe single-domain dataset, domain annotation requires a reference annotation method rather than human manual classification as the gold standard. The evaluation would then be uncontrollably biased towards methods that optimize for similar metrics. However, “good methods” under such metrics would make similar mistakes as reference methods. Also, false negatives of gold standard annotation methods can make methods with different optimization metrics (maybe more sensitive) produce a large number of high-scoring FPs.

### 4.7 Baselines

#### 4.7.1 Previously proposed methods

(1) Sequence search: MMseqs2 [9], BLASTp [10], HHblits [15, 16], EAT [44], and pLM-BLAST [46]. (2) Structure search — structural alphabet: 3D-BLAST-SW [21], CLE-SW [22], Foldseek, and Foldseek-TM [23]. (3) Structure search — structural alignment, reviewed in [60]: CE [24], Dali [25], and TM-align [26, 27]. For the specific introduction and settings of these proposed methods, see “Baseline details” Supplementary Section.

#### 4.7.2 Similarity-based search methods

These methods predict and sort the similarity between all query-target pairs (Fig. 5c). Different search methods are distinguished according to the way they predict similarity.

- Euclidean: Use the reciprocal of the Euclidean distance between embeddings.

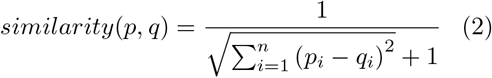
- COS: Use the COS distance between embeddings. *ϵ* is a small value to avoid division by zero.

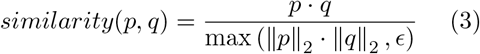
- SS-predictor: Combine the predicted TM-score with the top COS similarity. Unlike PLM-Search, which predicts pre-filtered pairs from PfamClan, SS-predictor extends its prediction to all protein pairs.

### 4.8 Experiment settings

#### 4.8.1 Pfam result generation

We obtained the Pfam family domains of proteins by PfamScan (version 1.6) [50] and Pfam dataset (Pfam36.0, 2023-09-12) [41]. For PfamClan, we query the comparison table Pfam-A.clans.tsv and replace the family domain with the clan domain it belongs to. For the family domain that has no corresponding clan domain, we treat it as a clan domain itself.

#### 4.8.2 Protein language model

ESMs are a set of protein language models that have been widely used in recent years. We used ESM-1b (650M parameters) [34], a SOTA general-purpose protein language model, to efficiently generate per-protein embeddings for PLMSearch.

For PLMAlign, the more extensive ProtT5-XL-UniRef50 (3B parameters) is used to generate per-residue embeddings. We conducted a detailed evaluation and analysis of the results from ESM-1b and ProtT5-XL-UniRef50 embeddings, as elaborated in “Remote homology alignment” Supplementary Section (Supplementary Fig. 7, Supplementary Table 8). Based on the analysis, ProtT5-XL-UniRef50 was selected.

#### 4.8.3 SS-predictor training

We used the deep learning framework PyTorch (version 1.7.1), ADAM optimizer, with MSE as the loss function to train SS-predictor. The batch size was 100, and the learning rate was 1e-6 on 200 epochs. The training ground truth was the TM-score calculated by TM-align. The datasets used for training (Extended Data Table A8) include: (1) All protein pairs from SCOPe40-train (8,953 proteins; 80,156,209 query-target pairs). (2) Undersampled protein pairs from CATHS40 (21,474 proteins; 28,440,312 query-target pairs). The parameters in ESM-1b were frozen and only the parameters in the bilinear projection network were trained.

#### 4.8.4 Experimental environment

We conducted the experiments on a server with a 56-core Intel(R) Xeon(R) CPU E5-2680 v4 @ 2.40 GHz and 256 GB RAM. The environment of our publicly available web server is 64 * Intel(R) Xeon(R) CPU E5-2682 v4 @ 2.50 GHz and 512 GB RAM.

## Supporting information

PLMSearch_supplement.pdf

## 5 Data availability

The sequences and structures of the SCOPe40 dataset are available at https://scop.berkeley.edu. The sequences of the Swiss-Prot dataset are freely available under the Creative Commons Attribution (CC BY 4.0) License at https://www.uniprot.org. The predicted structures are freely available from the AlphaFold Protein Structure Database at https://alphafold.ebi.ac.uk/download. CATH domain sequences and structures are publicly available at http://www.cathdb.info. Malidup can be found at http://prodata.swmed.edu/malidup; Malisam can be found at http://prodata.swmed.edu/malisam. Pfam is freely available under the Creative Commons Zero (‘CC0’) license at https://pfam.xfam.org. For PfamClan, we query the comparison table Pfam-A.clans.tsv at https://ftp.ebi.ac.uk/pub/databases/Pfam/current_release/Pfam-A.clans.tsv.gz. ESM-1b protein language model is available at https://github.com/facebookresearch/esm.

ProtT5-XL-UniRef50 protein language model is available at https://github.com/agemagician/ProtTrans. Source data of our work is provided at: (1) PLMSearch: https://dmiip.sjtu.edu.cn/PLMSearch/static/download/plmsearch_data.tar.gz. (2) PLMAlign: https://dmiip.sjtu.edu.cn/PLMAlign/static/download/plmalign_data.tar.gz.

## 6 Code availability

PLMSearch is freely available at https://dmiip.sjtu.edu.cn/PLMSearch. PLMAlign is freely available at https://dmiip.sjtu.edu.cn/PLMAlign. PLMSearch and related tutorials are freely available to the public at GitHub https://github.com/maovshao/PLMSearch/blob/main/pipeline.ipynb. Reproducing our results and regenerating the main and supplementary figures requires only one file at GitHub https://github.com/maovshao/PLMSearch/blob/main/main.ipynb. Run PLMAlign and reproduce the alignment experiment in “Remote homology alignment” Supplementary Section at GitHub: https://github.com/maovshao/PLMAlign.

## 7 Acknowledgements

This work has been supported by the National Natural Science Foundation of China (Grant No. 62272105), the Shanghai Municipal Science and Technology Major Project (Grant No. 2018SHZDZX01), the ZJ Lab, the Shanghai Research Center for Brain Science and Brain-inspired Intelligence Technology, and Beijing Academy of Artificial Intelligence (BAAI).

## 8 Author contributions

S.Z. conceived the project. S.Z. and J.Y. supervised the project. W.L., S.Z., and J.Y. designed the research and performed the analyses. W.L. wrote the software. H.W. sorted out the structure data. W.L. wrote the first draft of the manuscript. All authors contributed to the revision of the manuscript prior to submission and all authors read and approved the final version.

## 10 Figures and Tables

**Extended Data Fig. A1.**
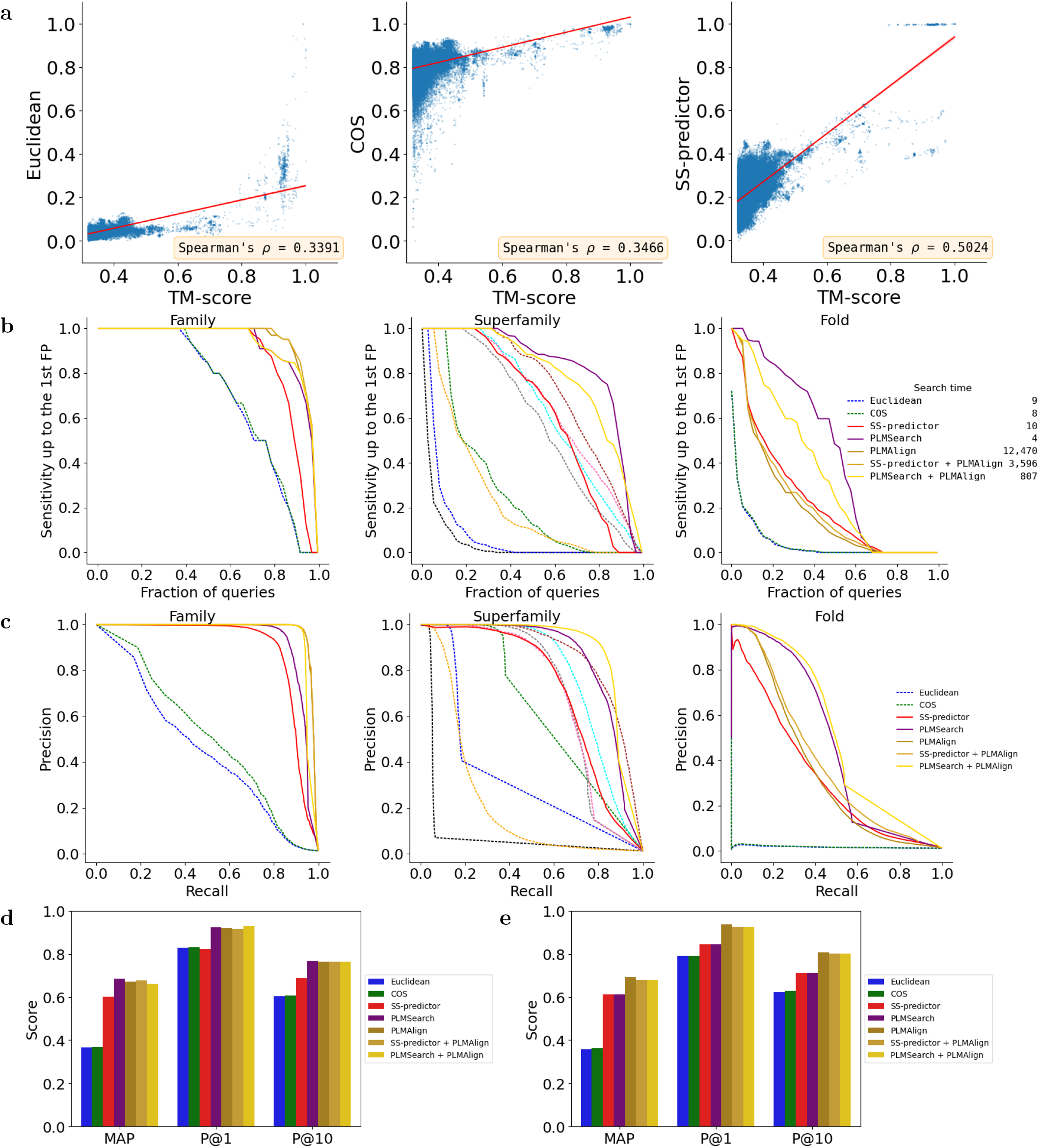
Ablation experiments: PfamClan, SS-predictor, and PLMAlign make PLMSearch more robust. **a**, Two-dimensional scatter plot of the predicted similarity and TM-score. From left to right are Euclidean, COS, and SS-predictor. We selected 100,000 protein pairs with highest TM-scores from the search results of five queries (with Swiss-Prot as the target dataset, 100,000 among a total of 2,150,700 query-target pairs) and used Euclidean, COS, and SS-predictor as the predicted similarity. We normalized the predicted similarity to 0-1 as the y-axis and their TM-scores (between 0-1) as the x-axis, thereby plotting the 100,000 protein pairs as points on a 2D plane. SS-predictor obtained the highest correlation coefficient with TM-score. **b-e**, Ablation experiments, with the same metrics used in Fig. 2. Extended Data Table A1 and Extended Data Table A3 record the specific values of each metric.

**Extended Data Fig. A2.**
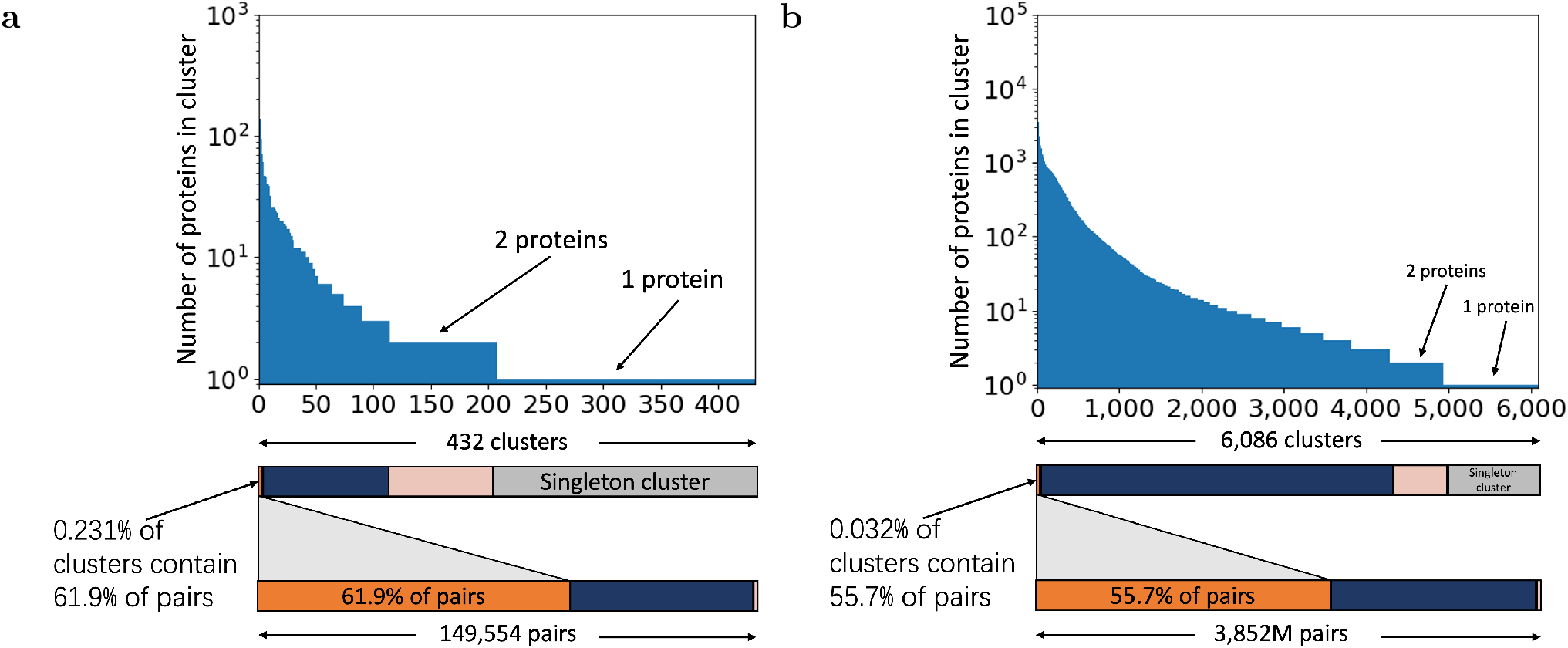
Clustering results based on Pfam clan on SCOPe40-test and Swiss-Prot. **a**, SCOPe40-test. **b**, Swiss-Prot. Proteins belonging to the same Pfam clan are clustered. The clustering results show a significant long-tailed distribution. After pre-filtering with PfamClan, more than 50% of the pre-filtered protein pairs (orange rectangles in the figure) are from the largest 1-2 clusters (big clusters), which only accounts for a very small part of the entire clusters (SCOPe40-test: 0.231%; Swiss-Prot: 0.032%). Therefore, big clusters will result in a significant number of irrelevant protein pairs in the pre-filtering results, reducing accuracy, and must be further sorted and filtered based on similarity, which is what SS-predictor does. See Supplementary Table 3 for specific statistical data.

**Extended Data Table A1.** All-versus-all search test on the SCOPe40-test dataset. The definition of AUROC, AUPR, MAP, and P@K is detailed in “Metrics” Section. The highest value achieved for each metric is highlighted in bold. Due to the width limit, Family and Superfamily are abbreviated as Fam and Sfam in the table, resepctively. The total search time spent for the all-versus-all search test is recorded.

| Methods | AUROC |  |  | AUPR |  |  | MAP | P@K |  | Time |
| --- | --- | --- | --- | --- | --- | --- | --- | --- | --- | --- |
|  | Fam | Sfam | Fold | Fam | Sfam | Fold | MAP | P@1 | P@10 | Seconds |
| Sequence search |  |  |  |  |  |  |  |  |  |  |
| MMseqs2 | 0.318 | 0.050 | 0.002 | 0.430 | 0.091 | 0.014 | 0.147 | 0.668 | 0.260 | **2 s** |
| Blastp | 0.527 | 0.161 | 0.004 | 0.717 | 0.342 | 0.029 | 0.183 | 0.717 | 0.354 | 10 s |
| HHblits | 0.920 | 0.363 | 0.064 | 0.969 | 0.623 | 0.256 | 0.320 | 0.858 | 0.577 | 10,998 s |
| EAT | 0.648 | 0.230 | 0.025 | 0.646 | 0.225 | 0.020 | 0.350 | 0.813 | 0.575 | 27 s |
| pLM-BLAST | 0.940 | 0.642 | 0.176 | 0.973 | 0.779 | 0.305 | 0.659 | 0.921 | 0.760 | 18,812 s |
| Structure search — structural alphabet |  |  |  |  |  |  |  |  |  |  |
| 3D-BLAST-SW | 0.653 | 0.255 | 0.045 | 0.621 | 0.264 | 0.047 | 0.446 | 0.825 | 0.604 | - |
| CLE-SW | 0.672 | 0.265 | 0.033 | 0.432 | 0.171 | 0.035 | 0.440 | 0.814 | 0.592 | - |
| Foldseek | 0.883 | 0.584 | 0.214 | 0.921 | 0.703 | 0.320 | 0.598 | 0.908 | 0.751 | 12 s |
| Foldseek-TM | 0.898 | 0.664 | 0.296 | 0.906 | 0.695 | 0.337 | 0.626 | 0.905 | 0.756 | 173 s |
| Structure search — structural alignment |  |  |  |  |  |  |  |  |  |  |
| CE | 0.847 | 0.527 | 0.148 | 0.882 | 0.627 | 0.245 | 0.618 | 0.897 | 0.734 | - |
| Dali | 0.923 | 0.702 | 0.281 | 0.948 | 0.814 | 0.454 | 0.702 | 0.927 | 0.790 | - |
| TM-align | 0.935 | 0.721 | 0.346 | 0.971 | 0.866 | **0.569** | **0.781** | **0.941** | **0.806** | 11,303 s |
| Our methods |  |  |  |  |  |  |  |  |  |  |
| Euclidean | 0.699 | 0.309 | 0.039 | 0.456 | 0.107 | 0.016 | 0.364 | 0.829 | 0.603 | 9 s |
| COS | 0.705 | 0.316 | 0.040 | 0.514 | 0.130 | 0.017 | 0.367 | 0.830 | 0.606 | 8 s |
| SS-predictor | 0.869 | 0.623 | 0.225 | 0.891 | 0.713 | 0.324 | 0.601 | 0.821 | 0.686 | 10 s |
| PLMSearch | 0.928 | **0.826** | **0.438** | 0.931 | 0.849 | 0.473 | 0.685 | 0.922 | 0.765 | 4 s |
| PLMAlign | 0.946 | 0.652 | 0.196 | 0.974 | 0.807 | 0.354 | 0.670 | 0.919 | 0.763 | 12,470 s |
| SS-predictor + PLMAlign | **0.949** | 0.665 | 0.211 | **0.975** | 0.822 | 0.391 | 0.677 | 0.915 | 0.763 | 3,596 s |
| PLMSearch + PLMAlign | 0.933 | 0.787 | 0.342 | 0.956 | **0.887** | 0.521 | 0.660 | 0.928 | 0.763 | 807 s |

**Extended Data Table A2.** The average number of family TPs, superfamily TPs, fold TPs, and total TPs up to the first FP on the SCOPe40-test search test. The average number of the total TPs also means the average rank of the first FP. The highest value achieved for each metric is highlighted in bold.

| Methods | Family | Superfamily | Fold | Total |
| --- | --- | --- | --- | --- |
| Sequence search |  |  |  |  |
| MMseqs2 | 2.20 | 0.51 | 0.02 | 2.74 |
| Blastp | 4.39 | 1.21 | 0.05 | 5.65 |
| HHblits | 9.10 | 11.25 | 3.92 | 24.29 |
| EAT | 6.17 | 6.12 | 1.65 | 13.96 |
| pLM-BLAST | 8.96 | 30.57 | 13.95 | 53.50 |
| Structure search — structural alphabet |  |  |  |  |
| 3D-BLAST-SW | 6.95 | 8.92 | 1.44 | 17.32 |
| CLE-SW | 6.97 | 9.65 | 1.10 | 17.74 |
| Foldseek | 8.57 | 28.33 | 14.19 | 51.10 |
| Foldseek-TM | 8.68 | 34.62 | 20.79 | 64.11 |
| Structure search — structural alignment |  |  |  |  |
| CE | 8.70 | 23.82 | 8.72 | 41.25 |
| Dali | 9.23 | 33.62 | 16.74 | 59.60 |
| TM-align | 9.29 | 38.57 | 24.83 | 72.70 |
| Our methods |  |  |  |  |
| Euclidean | 6.88 | 8.52 | 1.70 | 17.12 |
| COS | 6.94 | 8.79 | 1.77 | 17.52 |
| SS-predictor | 8.78 | 34.88 | 20.62 | 64.29 |
| PLMSearch | **9.32** | **47.44** | **48.01** | **104.78** |
| PLMAlign | 9.04 | 32.40 | 16.68 | 58.12 |
| SS-predictor + PLMAlign | 9.10 | 33.60 | 18.27 | 60.97 |
| PLMSearch + PLMAlign | 9.11 | 41.50 | 32.07 | 82.69 |

**Extended Data Table A3.** Evaluation on new proteins. See “New protein search test” Section. The definition of MAP, P@K is detailed in “Metrics” Section. The highest value achieved for each metric is highlighted in bold. The total search time spent for the search test is recorded.

|  | MAP | P@K |  | Time |
| --- | --- | --- | --- | --- |
| Methods | MAP | P@1 | P@10 | Seconds |
| Sequence search |  |  |  |  |
| MMseqs2 | 0.107 | 0.518 | 0.149 | **0.1 s** |
| Blastp | 0.132 | 0.590 | 0.270 | 0.5 s |
| HHblits | 0.279 | 0.872 | 0.581 | 548.1 s |
| EAT | 0.315 | 0.790 | 0.576 | 1.3 s |
| pLM-BLAST | 0.682 | 0.936 | 0.805 | 937.6 s |
| Structure search — structural alphabet |  |  |  |  |
| 3D-BLAST-SW | 0.383 | 0.763 | 0.580 | - |
| CLE-SW | 0.392 | 0.781 | 0.571 | - |
| Foldseek | 0.521 | 0.863 | 0.730 | 0.6 s |
| Foldseek-TM | 0.560 | 0.881 | 0.740 | 8.6 s |
| Structure search — structural alignment |  |  |  |  |
| CE | 0.580 | 0.845 | 0.722 | - |
| Dali | 0.643 | 0.909 | 0.804 | - |
| TM-align | **0.776** | **0.945** | **0.826** | 563.3 s |
| Our methods |  |  |  |  |
| Euclidean | 0.358 | 0.790 | 0.621 | 0.4 s |
| COS | 0.363 | 0.790 | 0.628 | 0.4 s |
| SS-predictor | 0.612 | 0.845 | 0.712 | 0.5 s |
| PLMSearch | 0.612 | 0.845 | 0.712 | 0.5 s |
| PLMAlign | 0.692 | 0.936 | 0.807 | 621.5 s |
| SS-predictor + PLMAlign | 0.679 | 0.927 | 0.801 | 179.2 s |
| PLMSearch + PLMAlign | 0.679 | 0.927 | 0.801 | 179.2 s |

**Extended Data Table A4.** Total running time of the web server. The environment of the web server is CPU ONLY, with 64 * Intel(R) Xeon(R) CPU E5-2682 v4 @ 2.50 GHz and 512 GB RAM. The time required to search 1, 10, and 100 query proteins with Swiss-Prot (568K proteins, the original dataset without filtering) and UniRef50 (53.6M proteins) as the target dataset were counted respectively.

| Query num | 1 | 10 | 100 |
| --- | --- | --- | --- |
| Swiss-Prot (568K proteins) |  |  |  |
| SS-predictor | 0.2 min | 0.5 min | 10.3 min |
| PLMSearch | 0.2 min | 1.1 min | 15.6 min |
| UniRef50 (53.6M proteins) |  |  |  |
| SS-predictor | 1.6 min | 6.3 min | 60.2 min |
| PLMSearch | 2.3 min | 12.1 min | 114.6 min |

**Extended Data Table A5.** The recall rate of different methods for easy pairs and remote homology pairs. We selected the 5000 pairs with the highest similarity for different search methods and counted the recalled and missed pairs. As shown in Fig. 3 **c-h**, “Easy pairs” refers to the protein pairs with similar sequences and similar structures in the first quadrant. “Remote homology pairs” refers to the protein pairs with dissimilar sequences but similar structures in the fourth quadrant. “All pairs” refers to all protein pairs with TM-score > 0.5 in the first and fourth quadrants.

| Methods | All pairs |  | Easy pairs |  | Remote homology pairs |  |
| --- | --- | --- | --- | --- | --- | --- |
|  | Recall | Miss | Recall | Miss | Recall | Miss |
| MMseqs2 | 0.450 | 0.549 | **1.000** | **0.000** | 0.165 | 0.834 |
| Blastp | 0.462 | 0.537 | **1.000** | **0.000** | 0.183 | 0.816 |
| Foldseek | 0.898 | 0.101 | **1.000** | **0.000** | 0.845 | 0.154 |
| Foldseek-TM | 0.901 | 0.098 | **1.000** | **0.000** | 0.850 | 0.149 |
| SS-predictor | 0.950 | 0.049 | **1.000** | **0.000** | 0.924 | 0.075 |
| PLMSearch | **0.989** | **0.010** | **1.000** | **0.000** | **0.983** | **0.016** |

**Extended Data Table A6.**
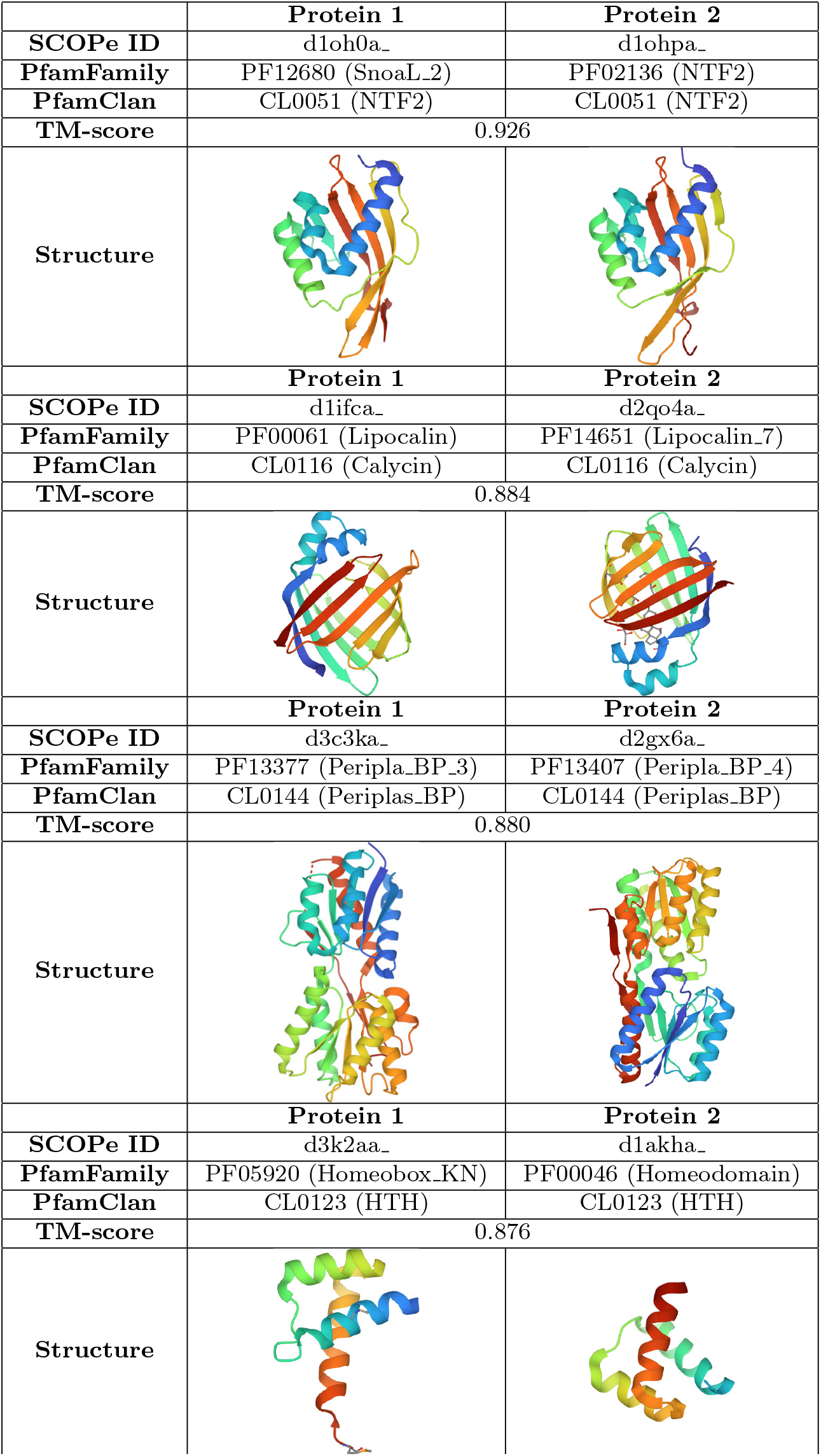
Case study for the pre-filtering results of PfamFamily & PfamClan. We investigated several protein pairs with TM-score>0.5 but missed by PfamFamily, and found that although the protein pairs do not share the same family, they contain similar families belonging to the same clan. Therefore, pre-filtering with PfamClan instead of Pfamfamily can help recall these protein pairs.

|  | Protein 1 | Protein 2 |
| --- | --- | --- |
| **SCOPe ID** | d1oh0a_ | d1ohpa_ |
| **PfamFamily** | PF12680 (SnoaL_2) | PF02136 (NTF2) |
| **PfamClan** | CL0051 (NTF2) | CL0051 (NTF2) |
| **TM-score** | 0.926 |  |
| **Structure** |  |  |
|  | Protein 1 | Protein 2 |
| **SCOPe ID** | d1ifca_ | d2qo4a_ |
| **PfamFamily** | PF00061 (Lipocalin) | PF14651 (Lipocalin_7) |
| **PfamClan** | CL0116 (Calycin) | CL0116 (Calycin) |
| **TM-score** | 0.884 |  |
| **Structure** |  |  |
|  | Protein 1 | Protein 2 |
| **SCOPe ID** | d3c3ka_ | d2gx6a_ |
| **PfamFamily** | PF13377 (Peripla_BP_3) | PF13407 (Peripla_BP_4) |
| **PfamClan** | CL0144 (Periplas_BP) | CL0144 (Periplas_BP) |
| **TM-score** | 0.880 |  |
| **Structure** |  |  |
|  | Protein 1 | Protein 2 |
| **SCOPe ID** | d3k2aa_ | d1akha_ |
| **PfamFamily** | PF05920 (Homeobox_KN) | PF00046 (Homeodomain) |
| **PfamClan** | CL0123 (HTH) | CL0123 (HTH) |
| **TM-score** | 0.876 |  |
| **Structure** |  |  |

**Extended Data Table A7.**
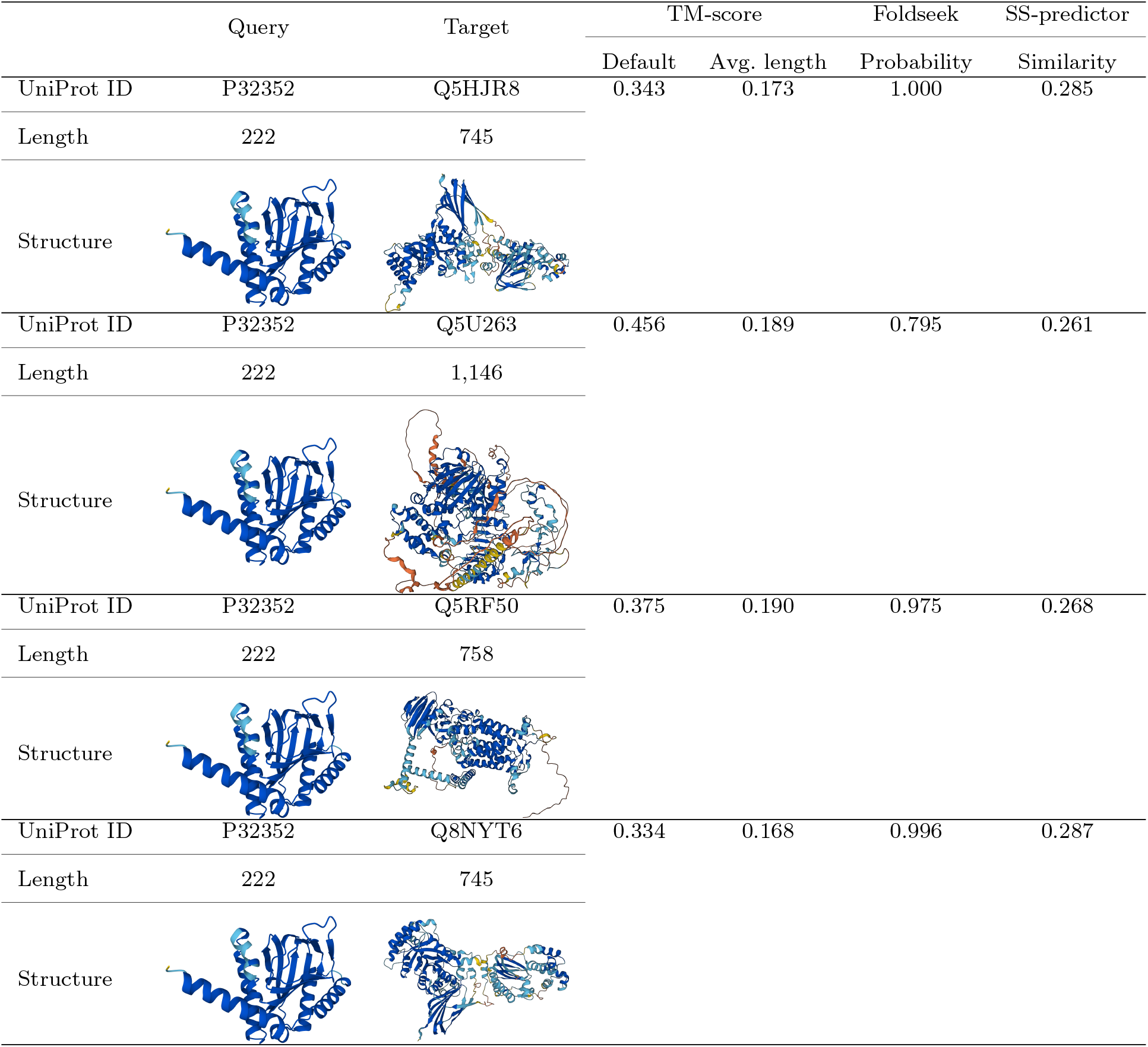
Four protein pairs selected for the manual inspection of protein pairs. They are filtered by Foldseek but with a TM-score<0.2 (Wrong pairs, defined in Fig. 3b). TM-align(Default) uses the query protein length as the normalized length. TM-align(Avg. length) uses the average length of protein pairs as the normalized length. As reported in Foldseek’s paper, Foldseek searches out these pairs because it focuses on local similarity. However, TM-align and PLMSearch focus on global similarity, so these pairs have TM-score<0.2 and similarity of SS-predictor lower than 0.3.

|  | Query | Target | TM-score |  | Foldseek | SS-predictor |
| --- | --- | --- | --- | --- | --- | --- |
|  |  |  | Default | Avg. length | Probability | Similarity |
| UniProt ID | P32352 | Q5HJR8 | 0.343 | 0.173 | 1.000 | 0.285 |
| Length | 222 | 745 |  |  |  |  |
| Structure |  |  |  |  |  |  |
| UniProt ID | P32352 | Q5U263 | 0.456 | 0.189 | 0.795 | 0.261 |
| Length | 222 | 1,146 |  |  |  |  |
| Structure |  |  |  |  |  |  |
| UniProt ID | P32352 | Q5RF50 | 0.375 | 0.190 | 0.975 | 0.268 |
| Length | 222 | 758 |  |  |  |  |
| Structure |  |  |  |  |  |  |
| UniProt ID | P32352 | Q8NYT6 | 0.334 | 0.168 | 0.996 | 0.287 |
| Length | 222 | 745 |  |  |  |  |
| Structure |  |  |  |  |  |  |

**Extended Data Table A8.** Datasets. By setting 0.4 sequence identity as the threshold to filter homologs, the max sequence identity of the test set relative to the training set does not exceed 0.4.

|  | Protein | Protein pair |
| --- | --- | --- |
| **Training** |  |  |
| **SCOPe40-train** | 8,953 | 80,156,209 (8,953 * 8,953)<br>TM-score>0.5: 504,553<br>TM-score≤0.5: 79,651,656 |
| **CATHS40** | 21,474 | 28,440,312 (undersampled)<br>TM-score>0.5: 7,813,946<br>TM-score≤0.5: 20,625,460 |
| **Test** |  |  |
| **SCOPe40-test** | 2,207 | 4,870,849 (2,207 * 2,207) |
| **New protein** | 110 | 242,770 (110 * 2,207) |
| **Swiss-Prot** | 430,140 | 43,014,000 (100 * 430,140) |
| **Target datasets on web server** |  |  |
| **Swiss-Prot (unfiltered)** | 568,744 | query num * 568,744 |
| **PDB** | 679,875 | query num * 679,875 |
| **UniRef50** | 53,625,855 | query num * 53,625,855 |
| **Evaluation on remote homology alignment** |  |  |
| **Malisam** | 233 | 129 |
| **Malidup** | 448 | 241 |

