## Supplementary material for "Protein language model powers accurate and fast sequence search for remote homology": PLMSearch_supplement.pdf

### 1 Supplementary Figures and Tables

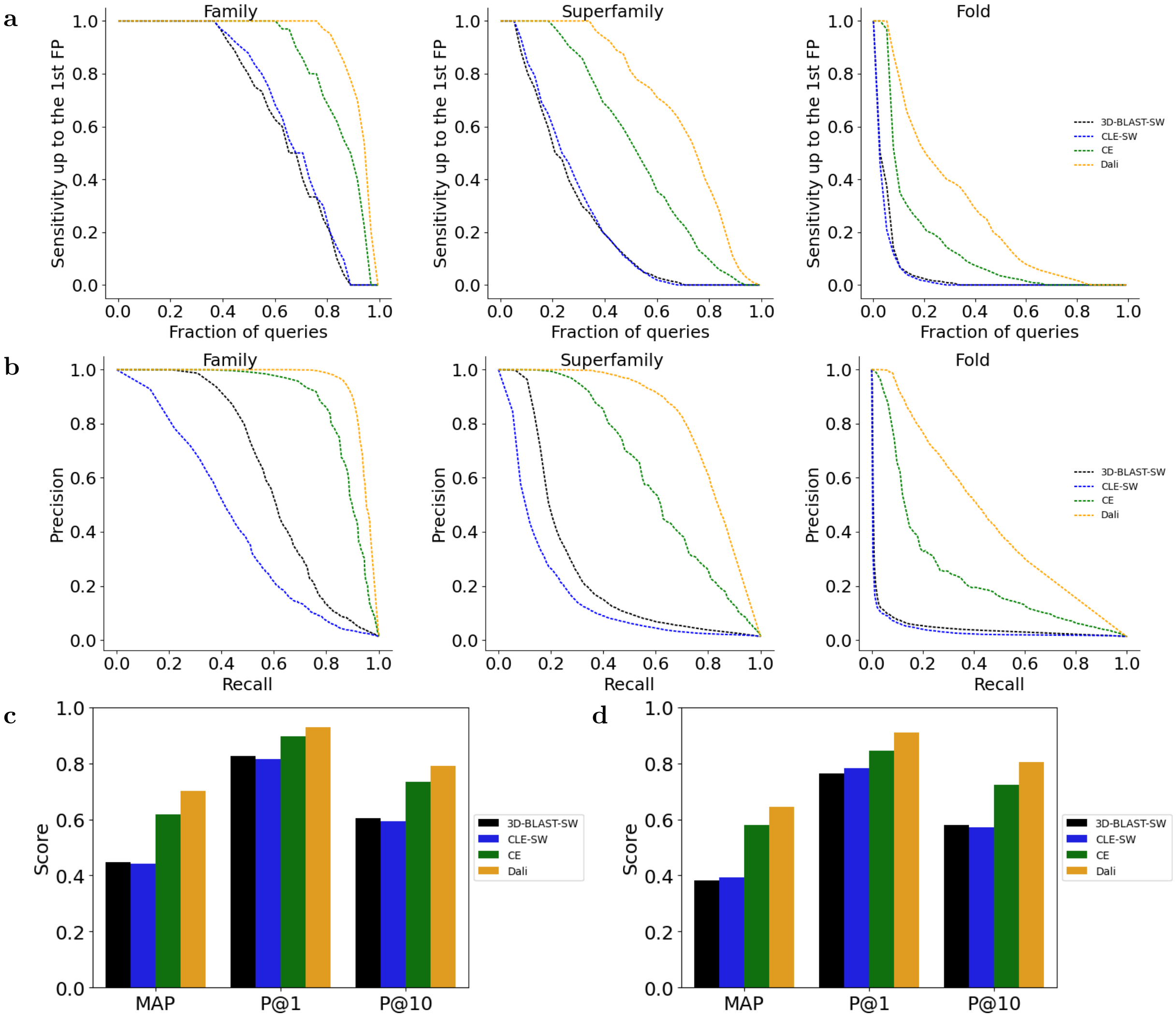


**Supplementary Fig. 1 Evaluation of other baselines.** The all-versus-all search test on SCOPe40-test with the same metrics as Fig. 2 in the main text. Extended Data Table A1 and Extended Data Table A3 record the specific values of each metric.

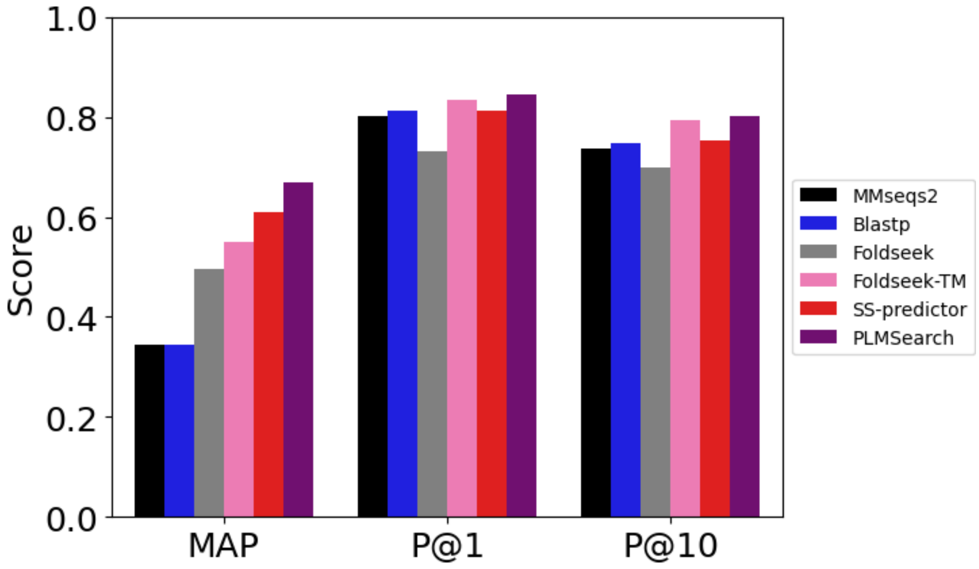


**Supplementary Fig. 2 MAP, P@1, and P@10 on the search test with Swiss-Prot as the target dataset.** Supplementary Table 2 records the specific values of each metric.

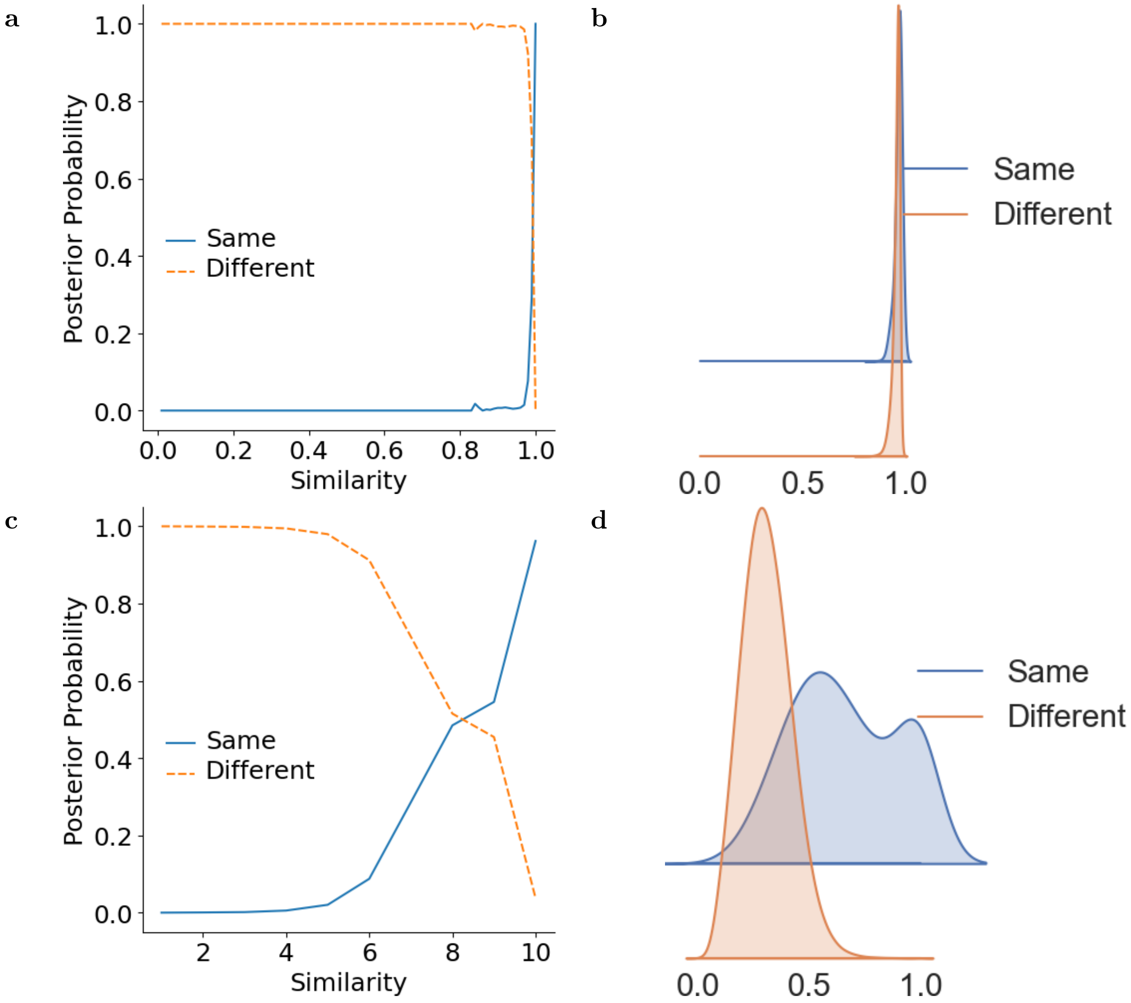


**Supplementary Fig. 3 Reference value of COS similarity and PLMAlign score. a-b**, COS similarity. **c-d**, PLMAlign score. **a, c** show the posterior probability of proteins with a given similarity being in the same fold or different folds in SCOPe40-train. **b, d** show the similarity distribution of the same fold and different folds protein pairs using kernel density estimation (smoothed histogram using a Gaussian kernel with the width automatically determined). The posterior probability corresponding to the similarity is shown in Supplementary Table 6. See "Reference similarity" Supplement Section for more details.

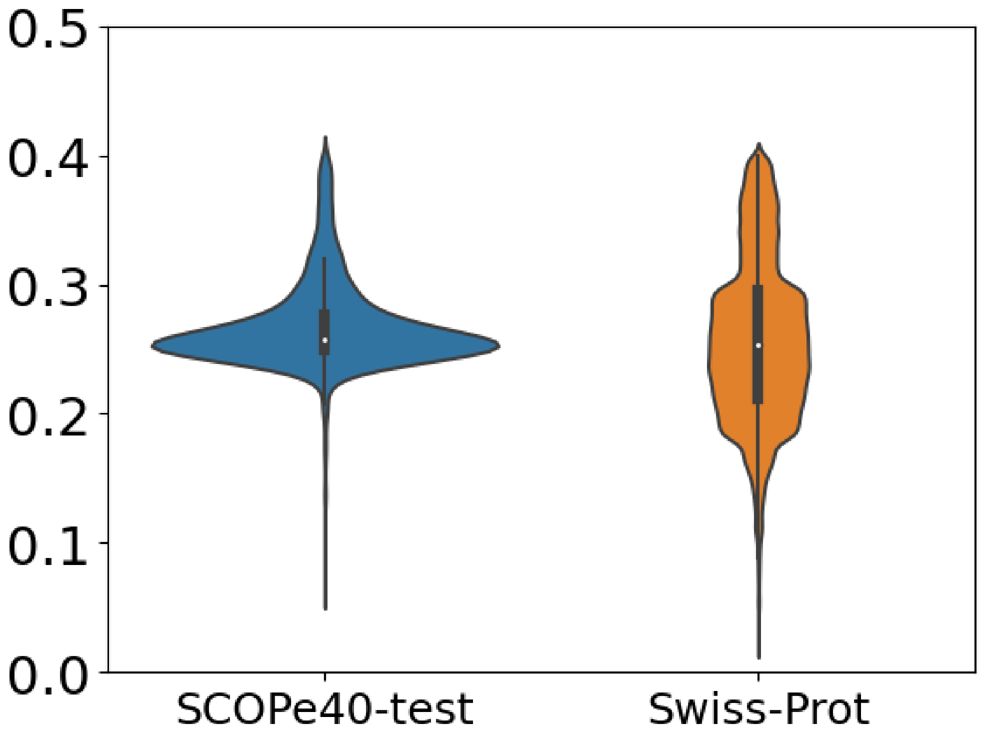


**Supplementary Fig. 4 The data distribution of max sequence identity of each protein in SCOPe40-test and Swiss-Prot against the training dataset (SCOPe40-train and CATHS40).** The majority of the maximum sequence identity is between 0.2 and 0.3. The sequence identity difference between their data is significantly bigger than that of pure random division, especially for the SCOPe40-test, which is the major test data, since the domains in SCOPe40-test belong to different folds with all domains in SCOPe40-train.

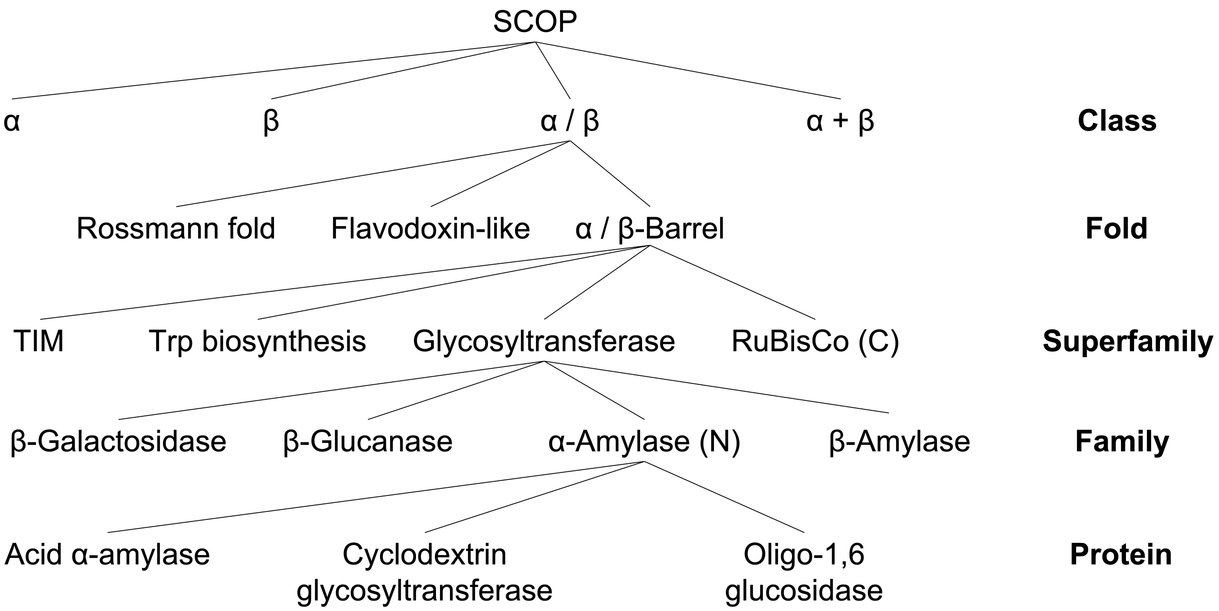


**Supplementary Fig. 5 Illustration of the SCOP hierarchy modified from Hubbard et al [1].**

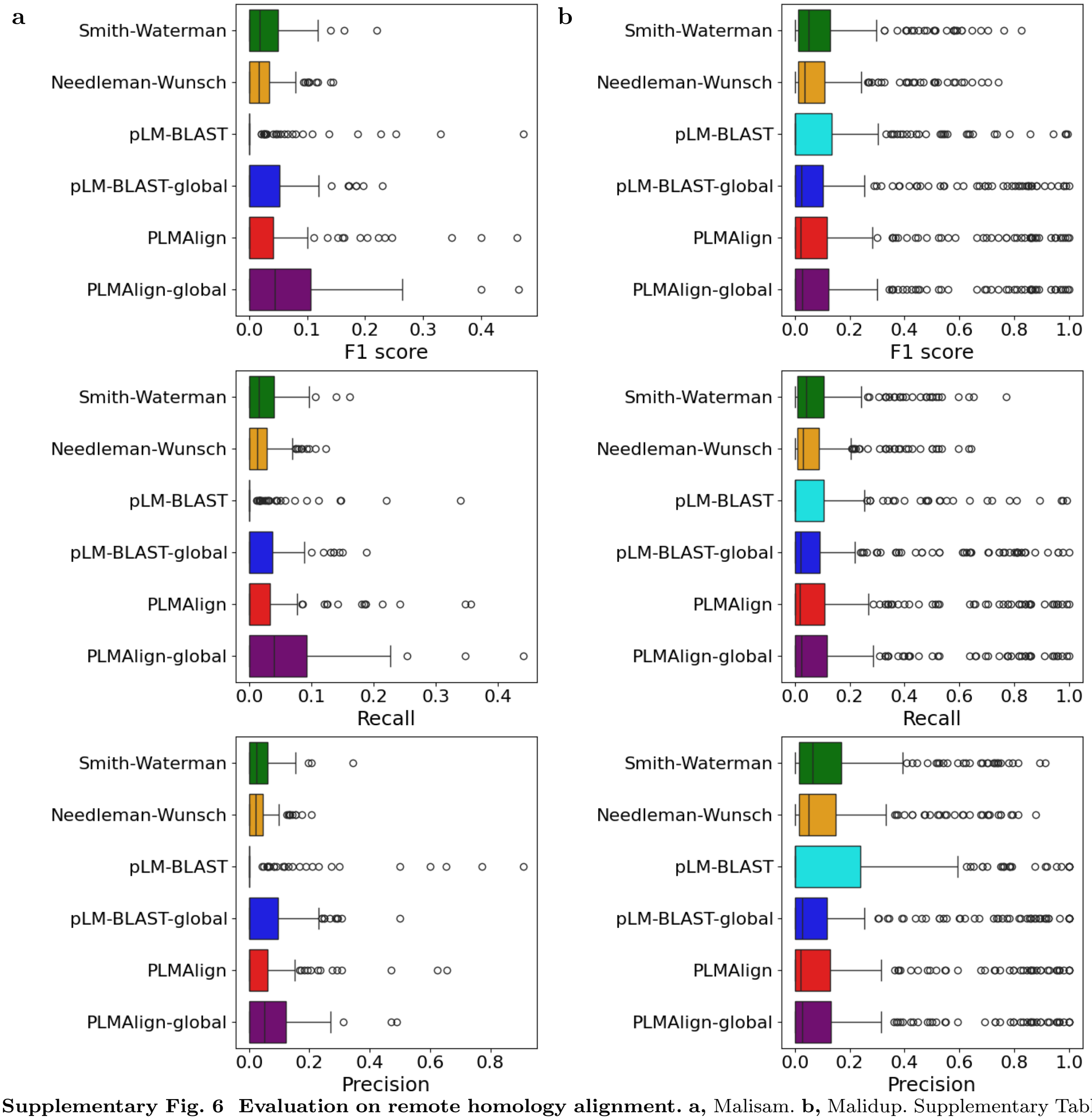


**Supplementary Fig. 6 Evaluation on remote homology alignment. a,** Malisam. **b,** Malidup. Supplementary Table 8 records the specific values of each metric.

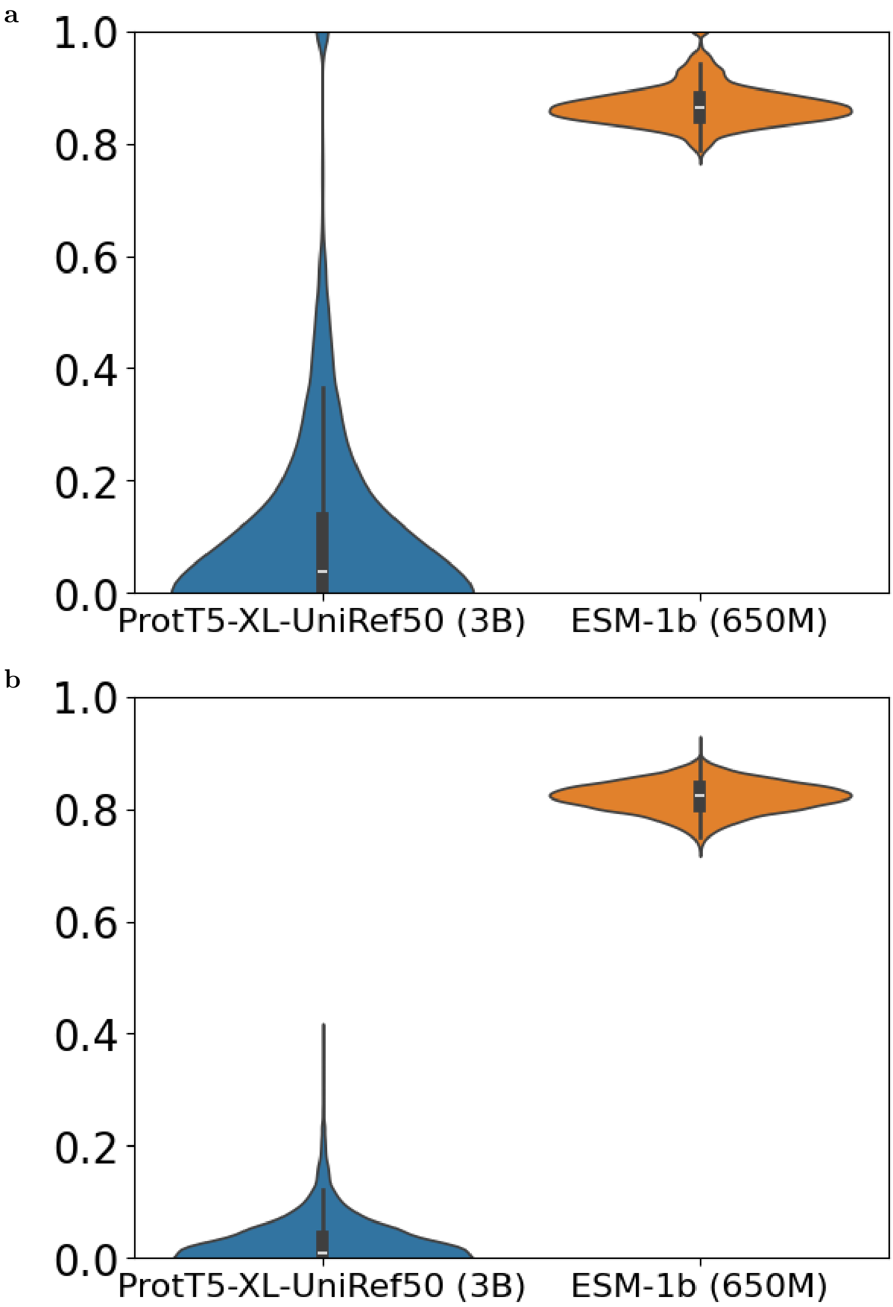


**Supplementary Fig. 7 Comparison of ESM-1b and ProtT5-XL-UniRef50.** The cos distance between the per-residue embeddings of two proteins. **a,** Self-alignment ($n*n$). **b,** Alignment with another protein ($n*m$). The COS distance between embeddings generated by ProtT5-XL-UniRef50 has greater coverage between 0 and 1, both in self-alignment and alignment with another protein.

| Methods | Search | | | | 5. Alignment | Total |
|---|---|---|---|---|---|---|
| | 1. Query embedding | 2. Query pfam | 3. Pfamclan | 4. SS-predictor | | |
| Swiss-Prot (568K proteins) | | | | | | |
| SS-predictor | 65 | 0 | 0 | 41 | 513 | 619 |
| PLMSearch | 65 | 35 | 29 | 28 | 779 | 936 |
| UniRef50 (53.6M proteins) | | | | | | |
| SS-predictor | 62 | 0 | 0 | 3,006 | 548 | 3,616 |
| PLMSearch | 62 | 38 | 3,324 | 2,893 | 563 | 6,880 |

**Supplementary Table 1 Running time of the web server at each step.** Search 100 query proteins with Swiss-Port (568K proteins) and UniRef50 (53.6M proteins) as the target dataset.

| | MAP | P@K | |
|---|---|---|---|
| Methods | MAP | P@1 | P@10 |
| Baselines | | | |
| MMseqs2 | 0.345 | 0.802 | 0.737 |
| Blastp | 0.343 | 0.812 | 0.748 |
| Foldseek | 0.497 | 0.732 | 0.697 |
| Foldseek-TM | 0.551 | 0.833 | 0.794 |
| Our methods | | | |
| SS-predictor | 0.610 | 0.812 | 0.754 |
| PLMSearch | **0.668** | **0.843** | **0.801** |

**Supplementary Table 2 Search test with Swiss-Prot as the target dataset.** TPs are protein pairs with TM-scores higher than 0.5. The definition of MAP and P@K is detailed in "Metrics" Section. The highest value achieved for each metric is highlighted in bold.

| **General statistics** | | | |
|---|---|---|---|
| Dataset | Protein num | Cluster num | Pair num |
| SCOPe40-test | 2,207 | 432 | 149,554 |
| Swiss-Prot | 430,140 | 6,086 | 3,852,993,796 |
| **Big cluster statistics** | | | |
| Dataset | Protein num | Cluster num | Pair num |
| SCOPe40-test | 305(13.8%) | 1(**0.231%**) | 92,720(**61.9%**) |
| Swiss-Prot | 65,453(15.2%) | 2(**0.032%**) | 2,149,740,012(**55.7%**) |
| **Small cluster statistics** | | | |
| Dataset | 2 proteins cluster num | 1 protein cluster num (Singleton cluster num) | |
| SCOPe40-test | 94 | 224 | |
| Swiss-Prot | 661 | 1,146 | |

**Supplementary Table 3 Statistics of clustering results based on Pfam clan on SCOPe40-test and Swiss-Prot.** The Big cluster in SCOPe40-test is CL0123. The Big clusters in Swiss-Prot are CL0023 and CL0063.

| Methods | Input | Sensitivity | Speed | Query mode |
|---|---|---|---|---|
| Sequence search | | | | |
| MMseqs2 | Sequence | Low | Very Fast | Multi query |
| Blastp | Sequence | Low | Very Fast | Multi query |
| HHblits | Profile HMMs | High | Slow | Single query |
| EAT | Per-protein embedding | Low | Very Fast | Multi query |
| pLM-BLAST | Per-residue embedding | Very High | Slow | Pairwise |
| Structure search — structural alphabet | | | | |
| Foldseek | Structure | High | Very Fast | Multi query |
| Foldseek-TM | Structure | Very High | Fast | Multi query |
| Structure search — structural alignment | | | | |
| TM-align | Structure | Very High | Slow | Pairwise |
| Our methods | | | | |
| SS-predictor | Per-protein embedding | High | Very Fast | Multi query |
| PLMSearch | Per-protein embedding | Very High | Very Fast | Multi query |
| PLMAlign | Per-residue embedding | Very High | Slow | Pairwise |

**Supplementary Table 4 Summary of the characteristics of baselines.** According to the performance on the all-versus-all search test on SCOPe40-test, the baselines are summarized according to their input, sensitivity, speed, and query mode.

| | **Search methods** | **Alignment methods** |
|---|---|---|
| **Input** | Per-protein embedding | Per-residue embedding |
| **Speed** | Very fast | Slow |
| **Similarity** | Yes | Yes |
| **How to obtain similarity** | Fast retrieval based on similarity prediction between embeddings | Pairwise alignment based on SW/NW, obtaining similarity from alignment scores |
| **Query mode** | Multi query | Pairwise |
| **Alignment (global or local)** | No | Yes |
| **Representation method** | PLMSearch, EAT | PLMAlign, pLM-BLAST |

**Supplementary Table 5 Differences between search methods and alignment methods.**

| SS-predictor | | | | | |
|---|---|---|---|---|---|
| Similarity | 0.1 | **0.3** | 0.5 | 0.7 | 0.9 |
| Posterior probability(same fold) | 0.000 | **0.003** | 0.456 | 1.000 | 1.000 |
| Posterior probability(different folds) | 1.000 | **0.996** | 0.543 | 0.000 | 0.000 |
| COS | | | | | |
| COS | 0.991 | 0.993 | **0.995** | 0.997 | 0.999 |
| Posterior probability(same fold) | 0.327 | 0.444 | **0.717** | 1.000 | 1.000 |
| Posterior probability(different folds) | 0.672 | 0.555 | **0.282** | 0.000 | 0.000 |
| PLMAlign | | | | | |
| Score | 3.0 | 5.0 | 7.0 | 9.0 | **9.5** |
| Posterior probability(same fold) | 0.001 | 0.020 | 0.285 | 0.545 | **0.749** |
| Posterior probability(different folds) | 0.998 | 0.979 | 0.714 | 0.454 | **0.250** |

**Supplementary Table 6 Posterior probability of SS-predictor similarity, COS similarity, and PLMAlign score in SCOPe40-train.** For SS-predictor, protein pairs with a similarity lower than 0.3 are usually assumed as randomly selected irrelevant protein pairs. For COS, the reference similarity 0.995 is selected. For PLMAlign, the reference score 9.5 is selected. See "Reference similarity" Supplement Section for more details.

| | Substitution matrix | Gap penalty | Scoring matrix | Traceback | Time |
|---|---|---|---|---|---|
| Smith-Waterman | Fix | Affine: 10+0.5*(L-1) | Truncate to zero | Begin with the highest score, end when 0 is encountered | - |
| Needleman-Wunsch | Fix | Affine: 10+0.5*(L-1) | Can be negative | Begin with the lower right of the matrix, end at top left | - |
| pLM-BLAST | Cosine | 0 | Can be negative | Traverse from all sequence boundaries | 122,564 s |
| pLM-BLAST-global | Cosine | 0 | Can be negative | Begin with the lower right of the matrix, end at top left | 18,812 s |
| PLMAlign | Dot Product | Linear: 1 * L | Truncate to zero | Begin with the highest score, end when 0 is encountered | 12,796 s |
| PLMAlign-global | Dot Product | Linear: 1 * L | Can be negative | Begin with the lower right of the matrix, end at top left | 12,470 s |

**Supplementary Table 7 Differences between the Smith-Waterman/Needleman-Wunsch algorithm, pLM-BLAST, and PLMAlign.** The analysis was conducted in four steps: Substitution matrix, Gap penalty, Scoring matrix, and Traceback. The total alignment time spent for the all-versus-all search test on SCOPe40-test (4,870,849 pairs) is recorded. Smith-Waterman and Needleman-Wunsch take the implementation of EMBL-EBI (https://www.ebi.ac.uk) as an example.

| Malisam | Number detected | F1 | Recall | Precision |
|---|---|---|---|---|
| Sequence | | | | |
| BLAST | 2 | 0.000 ± 0.000 | 0.000 ± 0.000 | 0.000 ± 0.000 |
| HMMER | 3 | 0.000 ± 0.000 | 0.000 ± 0.000 | 0.000 ± 0.000 |
| Needleman-Wunsch | 129 | 0.024 ± 0.002 | 0.020 ± 0.002 | 0.032 ± 0.003 |
| Smith-Waterman | 129 | 0.031 ± 0.003 | 0.025 ± 0.002 | 0.041 ± 0.004 |
| ProtT5-XL-UniRef50 | | | | |
| pLM-BLAST | 129 | 0.041 ± 0.008 | 0.031 ± 0.006 | **0.077 ± 0.014** |
| pLM-BLAST-global | 129 | 0.061 ± 0.009 | 0.053 ± 0.008 | 0.074 ± 0.011 |
| PLMAlign | 129 | 0.061 ± 0.010 | 0.059 ± 0.010 | 0.064 ± 0.010 |
| PLMAlign-global | 129 | **0.072 ± 0.010** | **0.069 ± 0.010** | 0.075 ± 0.010 |
| ESM-1b | | | | |
| pLM-BLAST | 129 | 0.020 ± 0.005 | 0.013 ± 0.003 | 0.047 ± 0.012 |
| pLM-BLAST-global | 129 | 0.032 ± 0.004 | 0.023 ± 0.003 | 0.058 ± 0.007 |
| PLMAlign | 129 | 0.037 ± 0.006 | 0.030 ± 0.005 | 0.050 ± 0.009 |
| PLMAlign-global | 129 | 0.068 ± 0.007 | 0.063 ± 0.006 | 0.076 ± 0.007 |

| Malidup | Number detected | F1 | Recall | Precision |
|---|---|---|---|---|
| Sequence | | | | |
| BLAST | 5 | 0.013 ± 0.013 | 0.006 ± 0.006 | 0.200 ± 0.200 |
| HMMER | 8 | 0.038 ± 0.026 | 0.021 ± 0.014 | 0.234 ± 0.153 |
| Needleman-Wunsch | 241 | 0.100 ± 0.009 | 0.081 ± 0.008 | 0.131 ± 0.012 |
| Smith-Waterman | 241 | 0.114 ± 0.010 | 0.094 ± 0.009 | 0.148 ± 0.013 |
| ProtT5-XL-UniRef50 | | | | |
| pLM-BLAST | 241 | 0.108 ± 0.013 | 0.091 ± 0.012 | 0.159 ± 0.017 |
| pLM-BLAST-global | 241 | 0.152 ± 0.017 | 0.139 ± 0.016 | **0.170 ± 0.019** |
| PLMAlign | 241 | 0.152 ± 0.017 | 0.147 ± 0.017 | 0.159 ± 0.018 |
| PLMAlign-global | 241 | **0.157 ± 0.017** | **0.151 ± 0.017** | 0.163 ± 0.018 |
| ESM-1b | | | | |
| pLM-BLAST | 241 | 0.106 ± 0.013 | 0.100 ± 0.013 | 0.141 ± 0.016 |
| pLM-BLAST-global | 241 | 0.133 ± 0.016 | 0.115 ± 0.014 | 0.164 ± 0.018 |
| PLMAlign | 241 | 0.129 ± 0.016 | 0.122 ± 0.015 | 0.140 ± 0.017 |
| PLMAlign-global | 241 | 0.151 ± 0.016 | 0.144 ± 0.015 | 0.159 ± 0.016 |

**Supplementary Table 8 Evaluation on remote homology alignment.** F1, Recall, and Precision are counted based on whether the generated alignment and manual alignment are consistent at each position. The highest value achieved for each metric is highlighted in bold.

| Methods | Similarity | Version |
|---|---|---|
| Sequence search | | |
| MMseqs2 | Bit score | Version 14.7e284 |
| Blastp | Bit score | Version 2.12.0+ |
| HHblits | Probability | Version 3.3.0 |
| EAT | 1 / (Embedding distance + 1) | Commit bcb935b |
| pLM-BLAST | Global similarity | Commit 0f226b0 |
| Structure search — structural alphabet | | |
| 3D-BLAST-SW | E-value in ascending order | Beta102, with BLAST+ 2.2.26 and SSW version ad452e |
| CLE-SW | Score | PDB Tool v4.80, SSW commit ad452e |
| Foldseek | Probability | Version 6.29e2557 |
| Foldseek-TM | Probability | Version 6.29e2557 |
| Structure search — structural alignment | | |
| CE | Z-score | BioJava's version 5.4.0 |
| Dali | Dali's Z-score | DaliLite.v5 |
| TM-align | TM-score | Version 20170708 |

**Supplementary Table 9 Similarity and versions of baselines.**

| Methods | Source |
|---|---|
| Sequence search | |
| MMseqs2 | https://github.com/soedinglab/MMseqs2 |
| Blastp | https://anaconda.org/bioconda/blast |
| HHblits | https://github.com/soedinglab/hh-suite |
| EAT | https://github.com/Rostlab/EAT |
| pLM-BLAST | https://github.com/labstructbioinf/pLM-BLAST |
| Structure search — structural alphabet | |
| 3D-BLAST-SW | http://3d-blast.life.nctu.edu.tw |
| CLE-SW | https://github.com/realbigws/PDB_Tool |
| Foldseek | https://github.com/steineggerlab/foldseek |
| Foldseek-TM | https://github.com/steineggerlab/foldseek |
| Structure search — structural alignment | |
| CE | https://github.com/biojava/biojava |
| Dali | http://ekhidna2.biocenter.helsinki.fi/dali |
| TM-align | https://seq2fun.dcmb.med.umich.edu/TM-align |

**Supplementary Table 10 Sources of baselines.**

| Methods | Family | Superfamily | Fold |
|---|---|---|---|
| MMseqs2 | | | |
| MMseqs2(Default) | 0.157 | 0.021 | 0.000 |
| MMseqs2(Best) | **0.318** | **0.050** | **0.002** |
| Foldseek | | | |
| Foldseek(Default) | 0.883 | 0.584 | 0.213 |
| Foldseek(Best) | 0.883 | 0.584 | 0.214 |
| Foldseek-TM(Best) | **0.898** | **0.664** | **0.296** |
| TM-align | | | |
| TM-align(Default) | 0.859 | 0.529 | 0.158 |
| TM-align(Avg. score) | 0.933 | 0.711 | 0.326 |
| TM-align(Avg. length) | **0.935** | **0.721** | **0.346** |

**Supplementary Table 11 Results with different settings for MMseqs2, Foldseek, and TM-align.** Different settings can greatly affect sensitivity. MMseqs2(Default) and Foldseek(Default) are the default settings of the program. MMseqs2(Best), Foldseek(Best), and Foldseek-TM(Best) are the practiced parameters in the experiments of Foldseek [2]. TM-align(Default) uses the query protein length as the normalized length. TM-align(Avg. score) calculates TM-scores for both comparison directions and averages them together. TM-align(Avg. length) uses the average length of protein pairs as the normalized length. We experimented with the settings that yielded the highest sensitivity, both as a metric and as a method. The results and setting are basically consistent with the conclusions obtained from Foldseek [2] and MT-LSTM [3].

### 2 Supplementary Note

#### 2.1 Sequence alignment

We used the same setting as BLAST programs. It could reflect the percentage of identical residues in the aligned sequence pairs. Sequence identity = (number of matched residues) / (the whole length of aligned sequences) [4]. We use the dynamic programming algorithm to perform pairwise sequence alignment and obtain the highest sequence identity among all alignments as the final sequence identity. The best alignment is also output as the sequence alignment result.

#### 2.2 Reference similarity

Researchers often want to know what similarity approximately corresponds to the protein pairs sharing the same fold. Here, we address this issue by calculating the posterior probability for proteins at certain similarities sharing the same or different folds. We will examine the results of the posterior probabilities using the fold standards defined by SCOP and protein pairs sharing the same fold are TPs. The experiments are performed with ramdomly selected 200 proteins from SCOPe40-train as queries and all proteins from SCOPe40-train as targets.

According to the Bayesian rules, for a given similarity, the posterior probabilities of proteins sharing the same or different folds can be expressed as:

$$\begin{cases} P(F \mid \mathrm{S}) = \frac{P(\mathrm{S}|F)P(F)}{P(\mathrm{S}|F)P(F)+P(\mathrm{S}|\bar{F})P(\bar{F})} \\ P(\bar{F} \mid \mathrm{S}) = \frac{P(\mathrm{S}|\bar{F})P(\bar{F})}{P(\mathrm{S}|F)P(F)+P(\mathrm{S}|\bar{F})P(\bar{F})} \end{cases} \tag{1}$$

Here, S stands for the similarity calculated by PLMSearch; $F$ and $\bar{F}$ represent the events that the protein pair shares the same and different folds in SCOP, respectively; $P(F)$ and $P(\bar{F})$ are the prior probabilities. $P(\mathrm{S} \mid F)$ and $P(\mathrm{S} \mid \bar{F})$ are the conditional probabilities of similarity when the two proteins are sharing the same or different folds, respectively. Thus, the conditional probabilities can be calculated by

$$\begin{cases} P(\mathrm{S} \mid F) = \frac{N(\mathrm{S})}{\sum N(\mathrm{S})} \\ P(\mathrm{S} \mid \bar{F}) = \frac{\bar{N}(\mathrm{S})}{\sum \bar{N}(\mathrm{S})} \end{cases} \tag{2}$$

where $N(S)$ is the number of protein pairs in the same fold with a certain similarity $S$, and $\bar{N}(S)$ is the number of protein pairs in the different folds with the similarity. The denominators are the summation of the same and different folds protein pairs.

The prior probabilities $P(F)$ and $P(\bar{F})$ can be calculated by

$$\begin{cases} P(F) = \frac{N(F)}{N(F)+N(\bar{F})} \\ P(\bar{F}) = 1 - P(F) \end{cases} \quad (3)$$

where $P(F)$ and $P(\bar{F})$ are, respectively, the numbers of the same and different folds pairs according to the SCOP definition. Overall, $P(F) = 0.0104$ and $P(\bar{F}) = 0.9896$ in our counting.

The posterior probability for two proteins with a certain similarity to be in the same SCOP Fold is calculated by integrating the data of Equations 2 and 3 into Equation 1.

#### 2.3 Remote homology alignment

##### 2.3.1 PLMAlign pipeline

The procedure of PLMAlign, akin to the Smith-Waterman and Needleman-Wunsch algorithm, primarily encompasses the following three steps:

- Calculation of the substitution matrix — Use dot product to replace the fixed values in the original substitution matrix.

  For a query protein of length $m$ and a target protein of length $n$, the per-residue embeddings are $E_m(m * d)$ and $E_n(n * d)$ respectively. The corresponding substitution matrix $S_{mn}(m * n)$ is then obtained by the cross product of these two matrices.

  $$S_{mn} = E_m \times E_n^T \quad (4)$$

  The essence of the cross product of the two matrices is that for the similarity $S_{mn}[i][j]$ between the i-th residue of the query protein and the j-th residue of the target protein, $S_{mn}[i][j]$ is calculated by the dot product of $E_m[i]$ and $E_n^T[j]$.

  $$S_{mn}[i][j] = E_m[i] \cdot E_n^T[j] \quad (5)$$

where $1 <= i <= m$ and $1 <= j <= n$. By replacing the fixed values in the original substitution matrix with the similarity (dot product) between vectors, PLMAlign is able to capture the evolutionary information in the context of residues and generates customized substitution matrices for each different query-target protein pair, resulting in more accurate alignments.

- Calculate the scoring matrix based on the substitution matrix and gap penalty — Linear gap penalty

  A linear gap penalty has the same scores for opening and extending a gap.

$$W_k = kW_1 \tag{6}$$

where $W_1$ is the cost of a single gap. The gap penalty is directly proportional to the gap length. When linear gap penalty is used, the Smith–Waterman algorithm can be simplified to:

$$H_{ij} = \max \begin{cases} H_{i-1,j-1} + s(a_i, b_j) \\ H_{i-1,j} - W_1 \\ H_{i,j-1} - W_1 \\ 0 \end{cases} \tag{7}$$

Compared with the traditional SW or NW algorithm using affine gap penalty (such as SW or NW implemented by EMBL-EBI (https://www.ebi.ac.uk)), the simplified algorithm uses $O(mn)$ steps, $m$ and $n$ are the lengths of the two sequences respectively. When an element is being scored, only the gap penalties from the elements that are directly adjacent to this element need to be considered, which greatly speeds up PLMAlign (Supplementary Table 7).

When performing global comparison, the score can be negative, and the corresponding score matrix calculation formula is:

$$H_{ij} = \max \begin{cases} H_{i-1,j-1} + s(a_i, b_j) \\ H_{i-1,j} - W_1 \\ H_{i,j-1} - W_1 \end{cases} \tag{8}$$

- Search path based on scoring matrix — Same as traditional SW or NW algorithm.

  When performing a local comparison, PLMAlign begins with the highest score, end when 0 is encountered. When performing a global comparison, PLMAlign begins with the cell at the lower right of the matrix, end at top left cell.

The differences between the SW/NW algorithm, pLM-BLAST, and PLMAlign are discussed in further detail in Supplementary Table 7.

##### 2.3.2 Evaluation on remote homology alignment

Manual structure alignment is an intuitive human assessment, typically emphasizing 3D overlap and topology preservation, as these features are easier to visualize than a multitude of local alignments and contacts [5, 6]. All methods tend to concur when the sequence identity is high. Therefore, the most valuable gold-standard alignment benchmark set is one where the dataset members exhibit low sequence identity and varied degrees of structural similarity. Our benchmarks were conducted on the curated Malisam [7] and Malidup [8] protein structural alignment benchmarking datasets, which are heavily skewed towards difficult-to-detect, low-sequence-identity remote homology.

As depicted in Supplementary Fig. 6 and Supplementary Table 8, in both benchmarks, the majority of the protein alignments failed to pass the filtering steps in both BLAST and HMMER. In other words, BLAST and HMMER were unable to detect the vast majority of the alignments. This left Smith–Waterman and Needleman–Wunsch as the baselines for sequence alignment methods. Owing to the use of dot products to calculate similarity instead of fixed values in the original substitution matrix, PLMAlign outperforms Smith–Waterman and Needleman–Wunsch. Moreover, compared to pLM-BLAST, PLMAlign performs better on F1 and Recall, possibly because PLMAlign taking the gap penalty into account. However, pLM-BLAST exhibits higher precision, possibly because pLM-BLAST employs traversal and take maximum value for path search. Through time comparison (Supplementary Table 7), we discovered that PLMAlign is faster, particularly in local alignment. This may be primarily due to: (1) Dot product is faster than Cosine as no normalization is required. (2) PLMAlign uses a linear gap penalty model. When considering the gap penalty for a certain position, only the adjacent upper and left positions need to be considered (without considering the entire column and row). (3) For local alignment only, PLMAlign directly searches from the maximum value of the entire matrix, rather than searching in a traversal manner.

Additionally, we explored the impact of different language model embeddings (Supplementary Fig. 7). We compared the per-residue embeddings generated by ESM-1b and ProtT5-XL-UniRef50. We found that the COS distance between embeddings generated by ProtT5-XL-UniRef50 has greater coverage between 0 and 1, both in self-alignment and alignment with another protein. This suggests that the embedding generated by ProtT5-XL-UniRef50 can more distinctly distinguish the similarity between residues. Through experimental verification, we also found that ProtT5-XL-UniRef50 can yield better alignment results (either for PLMAlign or pLM-BLAST, see Supplementary Table 8).

#### 2.4 Baseline details

We first summarize the similarity for sorting and versions of different methods in Supplementary Table 9, and summarize the sources of different methods in Supplementary Table 10.

##### 2.4.1 Sequence search

- MMseqs2: A sequence search method with huge improvements in speed and sensitivity over other sequence search methods. For MMseqs2, different parameter settings will bring huge differences in search results and have a huge impact on the sensitivity of search results. The default parameters (MMseqs2(Default)) lead to lower sensitivity Supplementary Table 11. In order to ensure the fairness of the experiment, we used the parameters practiced in the Foldseek paper (–threads 56 -s 7.5 -e 10000 –max-seqs 2000) for experiments (MMseqs2(Best)). MMseqs2 sorts the results by bit score as similarity.
- Blastp: We first downloaded Blastp from Anaconda with the order: conda install -c bioconda blast. Then, we used the default parameters to build target datasets for SCOPe40-test and Swiss-Prot and searched against them. Taking the SCOPe40-test as an example, the command to build the dataset: makeblastdb -in protein.fasta -title scope40 -dbtype prot -out scope40 -parse_seqids. Search command: blastp -query protein.fasta -db scope40 -out search_result -outfmt "6 qacc sacc bitscore" -num_threads 56.
- HHblits: We first downloaded HHblits from Anaconda with the order: conda install -c conda-forge -c bioconda hhsuite. Then, we used the default parameters to build target datasets for SCOPe40-test and search against it. 1. Download the UniRef30 database: wget https://gwdu111.gwdg.de/ compbiol/uniclust/2023_02/UniRef30_2023_02_hhsuite.tar.gz. 2. Build the SCOPe40-test dataset according to the series of commands in "Building customized databases" in wiki tutorial: https://github.com/soedinglab/hh-suite/wiki and search against it with hhblits.
- EAT: We complete the following steps according to a series of commands in the repository: https://github.com/Rostlab/EAT. 1. Install 2. Use ProtT5-XL-U50 (or ProtT5 for short) to calculate the embedding of each residue (Lx1024 for ProtT5). The embeddings for each protein are derived by

averaging the embeddings for each residue, resulting in a single 1024-d vector for each protein, regardless of its length, and the embeddings are stored as H5 files. 3. Calculate the inter-embedding Euclidean distance and sort according to 1 / (Embedding distance + 1) to complete the search.

- pLM-BLAST: We complete the following steps according to a series of commands in the repository: https://github.com/labstructbioinf/pLM-BLAST. 1. Install. 2. Use scripts/makeindex.py to generate index files from FASTA files. 3. Use the embeddings.py script to create the database. 4. Use dbtofile.py to create an additional file with flattened embeddings. 5. Use pLM-BLAST to search based on the generated embeddings.

##### 2.4.2 Structure search — structural alphabet

- 3D-BLAST-SW: We used 3D-BLAST (beta102) with BLAST+ (2.2.26) and SSW [9] (version ad452e). We first converted the PDB structures to a 3D-BLAST dataset using 3d-blast -sq_write and 3d-blast -sq_append. For Smith-Waterman we used (1) gap open of 8 (2) gap extend of 2 and (3) returning alignments (-c) (4) using the 3D-BLAST's optimized substitution matrix (-a 3DBLAST), (5) protein alignment mode (-p): ssw_test -o 8 -e 2 -c -a 3DBLAST -p. 3D-BLAST-SW's results are sorted by E-value in ascending order.
- CLE-SW: We used PDB Tool v4.80 (github.com/realbigws/PDB_Tool) to convert the benchmark structure set to CLE sequences. After the conversion, we used SSW (commit ad452e) to align CLE sequences all-versus-all. We sorted the results by alignment score. The following parameters were used to run SSW: (1) protein alignment mode (-p), (2) gap open penalty of 100 (-o 100), (3) gap extend penalty of 10 (-e 10), (4) CLE's optimized substitution matrix (-a cle.shen.mat), (5) returning alignment (-c). The gap open and extend values were inferred from DeepAlign [10]. The results are sorted by score in descending order. ssw_test -p -o 100 -e 10 -a cle.shen.mat -c.
- Foldseek & Foldseek-TM: The latest protein structure search method, which achieves extremely high sensitivity in protein search by directly using structural information for encoding. Similarly, differences in parameter settings also affect the sensitivity of the Foldseek method Supplementary Table 11. Again, we use the parameters practiced in the Foldseek paper (–threads 56 -s 9.5 -e 10 –max-seqs 2000) for experiments (FoldSeek(Best)). Foldseek-TM adds additional parameter "–alignment-type 1". Foldseek & Foldseek-TM sorts the results by probability.

##### 2.4.3 Structure search — structural alignment

- CE: We used BioJava's [11] (version 5.4.0) implementation of the combinatorial extension (CE) alignment algorithm. We modified one of the modules of BioJava under shape configuration to calculate the CE value. Our modified CEalign.jar file requires a list of query files, the path to the target PDB files, and an output path as input parameters. This Java module runs an all-versus-all CE calculation, with unlimited gap size (maxGapSize -1) to improve alignment results [12]. The results were sorted by Z-score in descending order. For the multi-domain benchmark, we excluded 1 query that was running over 16 days. The Jar file of our implementation of CE calculation is provided. java -jar CEalign.jar querylist.txt TargetPDBDirectory OutputDirectory.
- Dali: We installed the standalone DaliLite.v5. For the SCOPe40 benchmark set, input files were formatted in DAT files with Dali's import.pl. The conversion to DAT format produced 11,137 valid structures out of the 11,211 initial structures for the SCOPe benchmark, and 34,256 structures out of 34,270 spice clusters. After formatting the input files, we calculated the protein alignment with Dali's structural alignment algorithm. The results were sorted by Dali's Z-score in descending order: import.pl –pdbfile query.pdb –pdbid PDBid –dat DAT dali.pl –cd1 queryDATid –db targetDB.list –TITLE systematic –dat1 DAT –dat2 DAT –outfmt "summary" –clean.
- TM-align: We first downloaded TM-align from Anaconda with the order: conda install -c bioconda tmalign. We ran the benchmark using "-a" parameters. So TM-align reports three TM-scores: (1) normalized by the length of 1st chain (query), (2) normalized by the length of the 2nd chain (target), (3) normalized by the average length of two structures. TM-align(Avg. length) uses the TM-score normalized by the average length of two structures and outperforms other settings (Supplementary Table 11). So the TM-score used in this paper is generated by TM-align(Avg. length).
